# Api5 Regulates Genomic Stability and Chemotherapy Resistance in Cancer

**DOI:** 10.64898/2026.06.23.734059

**Authors:** Benchamin Abraham, Ayushi Upadhyay, Khushi Malhotra, Ajay J Malik, Dhananjay Virkar, Aryaman Deshmukh, Mayurika Lahiri

## Abstract

Api5 is elevated in a number of cancers and is associated with many hallmarks of cancer, including resistance to apoptosis, immune escape, stemness, chemotherapy resistance, high proliferation, and cell-cycle dysregulation. In this study, we identified the DNA and chromatin-binding activities of Api5 in tumorigenic cells, as well as its association with genomic instability and chemotherapy resistance. Knockdown of Api5 resulted in reduced nuclear volume, DNA content, and chromosome number, and increased sensitivity to DNA damage. The survival of Api5-knockdown cells decreased following UV and cisplatin treatments due to the accumulation of damaged DNA and inefficient nucleotide excision repair. Interestingly, Api5 knockdown cells also exhibited low pChk1 levels following UV damage. Further, we confirmed the chemotherapy resistance phenotype in cancers with elevated Api5 levels, demonstrating that xenograft tumours with Api5 knockdown responded better to cisplatin, with significant tumour regression.

**Summary:** Apoptosis inhibitor 5 (Api5) contributes to chemotherapy resistance by conferring a survival advantage and promoting efficient DNA repair following genotoxic stress through regulation of Chk1 activation.

## Introduction

Since the 1940s, clinicians have used chemotherapy to treat cancer by inducing cell death within tumour tissues using different cytotoxic compounds to achieve tumour remission. Among the numerous chemotherapeutic agents, compounds that induce apoptosis via DNA damage were developed earliest and are the most widely used. Cisplatin is one such alkylating agent that cross-links guanine at the N7 position, disrupting DNA replication and transcription and causing cell death (Nygren, 2001). Cancer cells resist chemotherapy by inhibiting apoptotic pathways and enhancing DNA repair pathways (Martin et al., 2008). The nucleotide excision repair (NER) pathway repairs DNA cross-links formed after alkylation and ultraviolet (UV) exposure (Basu and Krishnamurthy, 2010). The DDB1-DDB2 complex detects distortions caused by cross-linked nucleotides and initiates the sequential recruitment of NER proteins, including XPC, XPB, XPD, XPA, and XPG, to the damaged sites (Ray et al., 2013). Efficient repair of lesions requires activation of the ATR pathway, which ultimately activates Chk1, thereby promoting cell cycle arrest, giving cells time to repair DNA and preventing progression through the cell cycle with mutated DNA (Paulsen and Cimprich, 2007). Prolonged activation of the ATR pathway indicates unresolved DNA lesions, and in such conditions, Chk1 phosphorylates p53 to initiate apoptosis (Ou et al., 2005; Yazinski and Zou, 2016). Cancers become resistant to chemotherapy by altering the apoptotic regulators and DNA repair proteins, thus preventing apoptosis and increasing DNA repair (Bouwman and Jonkers, 2012; Pommier et al., 2004).

Apoptosis inhibitor 5 (Api5) is a multifunctional protein that is often overexpressed in many cancers (Cho et al., 2014; Koci et al., 2012; Sasaki et al., 2001). High levels of Api5 contribute to several key features of cancer, such as improved cell survival (Imre et al., 2017), increased cell growth (Garcia-Jove Navarro et al., 2013), stem-like properties (Bhatia and Lahiri, 2026; Song et al., 2017), immune evasion (Kim et al., 2018), the ability to spread to other tissues (Song et al., 2015), and resistance to chemotherapy drugs like cisplatin (Jang et al., 2017). Among these effects, Api5’s ability to prevent apoptosis (Morris et al., 2006) is most widely studied and is considered the main reason for its cancer-promoting activity.

Early studies showed that Api5 suppresses programmed cell death and exhibits high expression in several types of tumours, including B-cell lymphomas (Krejci et al., 2007), head and neck squamous cell carcinoma (Järvinen et al., 2008), cervical cancer (Cho et al., 2014), and other epithelial cancers (Basset et al., 2017; Bousquet et al., 2019). Api5 expression increases in the later stages of cancer, promoting cancer progression and poor patient survival (Cho et al., 2014; Kuttanamkuzhi et al., 2023). These observations led researchers to investigate how Api5 promotes cancer progression. Independent studies identified various functional pathways that could aid in cancer progression. One such noticeable study reported that Api5 prevents apoptosis by repressing E2F1-dependent transcription and inhibiting caspase-2 activation (Morris et al., 2006). Cancer cells have higher cell proliferation rates and Api5 enhances the proliferation too. E2F1-mediated cell cycle progression is significantly affected by Api5, where high Api5 levels drive the transition from the G1 phase to the S phase (Garcia-Jove Navarro et al., 2013). In addition to cell cycle regulation, Api5 promotes cancer stem-like properties by activating the FGF2–NANOG pathway and stimulates cell growth by interacting with oestrogen receptor-alpha (ER-α) (Basset et al., 2017). Although Api5 interacts with nuclear ER-α, it does not directly bind to ER-*α* target gene promoters, suggesting that its effects on gene expression occur through indirect mechanisms. Matrix metalloproteinases are proteinases secreted by metastatic cancers to degrade the basal membrane, thereby aiding invasion and metastasis (Egeblad and Werb, 2002). Elevated matrix metalloproteinase expression is a direct consequence of Api5-induced activation of the ERK pathway, which significantly enhances the invasive and metastatic potential of tumours (Song et al., 2015). In chemo-resistant triple-negative breast cancer, FGFR1 activation, leading to Bim degradation, plays a major role in poor treatment response of Api5-overexpressing cells (Jang et al., 2017).

Despite these findings, the role of Api5 in chemotherapy response and its high expression in the later stages of cancer remain poorly understood. Therefore, its possible role in maintaining genome stability remains a mystery. In this study, we discovered the DNA-binding and differential chromatin-binding ability of Api5. Using the MCF10 isogenic human breast cancer progression model (Bessette et al., 2015), we show, for the first time, that Api5 contributes to genomic instability by regulating Chk1 activation and influencing the NER pathway. This newly identified role shows a clear connection between high levels of Api5 and resistance to chemotherapy. Involvement of Api5 in Chk1 activation provides a new explanation for how cancer cells survive DNA damage induced by the chemotherapy drug cisplatin. By linking Api5 to important repair pathways that detect and repair DNA damage, this study improves our understanding of how Api5 contributes to cancer development. It also supports reports on targeting Api5 for new treatments, especially in cancers with genomic instability and chemotherapy resistance.

## Results

### Api5 is a chromatin-associated protein capable of directly binding to DNA

Api5 lacks a typical DNA-binding domain after its leucine zipper region (Figure 1A) (Han et al., 2012), yet the bioinformatics tool DRNApred predicted a potential DNA-binding ability of Api5. To show that Api5 binds to DNA, an electrophoretic mobility shift assay (EMSA) was performed, where a shift in the binding ability of Api5 to DNA increased with increasing concentrations of Api5 (Figure 1B).

**Figure 1:**
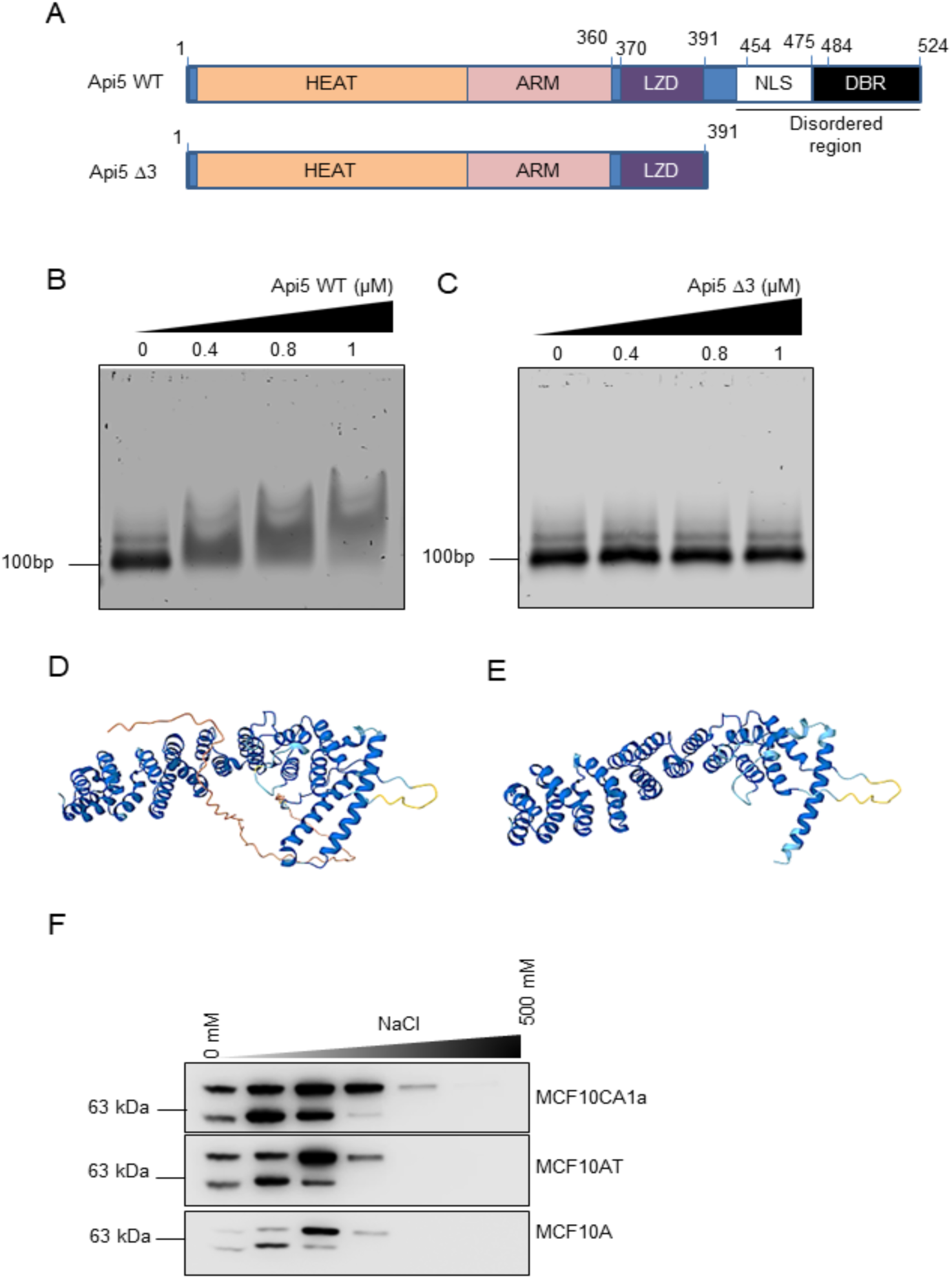
C-terminal region of Api5 is critical for DNA binding. **A.** Schematic representation of full-length Api5 and its deletion construct, Api5 Δ3. Api5 wild type (WT) protein is 524 amino acids long with HEAT repeats, ARM-like repeats at the N-terminal region that are known for protein-protein interactions, followed by leucin zipper domain (LZD), nuclear localisation signal (NLS) and DNA binding region (DBR). The predicted DBR is deleted in Api5 Δ3. **B**. EMSA was performed using 100ng poly dA:dT 100-mer DNA and increasing concentrations of full-length Api5. **C.** EMSA performed using 100ng poly dA:dT 100-mer DNA and increasing concentrations of Api5 Δ3. **D** and **E** are the alpha fold predicted structures of Api5 WT and Api5 Δ3 proteins. **F**. Sequential salt extraction (SSE) of chromatin-bound proteins performed in the MCF10 isogenic cell line series that includes non-malignant MCF10A, pre-malignant MCF10AT and malignant MCF10CA1a cells.

Further analysis of DNA-binding ability using the bioinformatic tool DP-Bind identified a DNA-binding region spanning amino acids 466–524 at the C-terminus (Table S1). Deletion of 133 amino acids from the C-terminal end, which included this predicted region, generated the Api5 Δ3 mutant (Figure 1A), which did not show any shift in the EMSA assay and therefore was not able to bind DNA (Figure 1C). However, the overall folding and structure of the mutant protein remained similar to wild-type Api5 (Api5 WT) (Figure 1D and E). The deletion of 40 amino acids from the C-terminus, which harbours multiple lysine and arginine residues, did not affect the DNA binding ability compared to the Api5 WT (Figure S1 A-D).

As Api5 has been reported to be a nuclear protein associated with chromatin (Garcia-Jove Navarro, 2013), we investigated its chromatin-binding ability in the isogenic MCF10 breast cancer progression model. Within this isogenic series, MCF10A, MCF10AT, and MCF10CA1a cells represent non-tumorigenic, premalignant, and malignant phenotypes, respectively. Api5 expression increased in proportion to malignancy across the cell line series, consistent with previous reports in which Api5 levels increase across different cancer stages (Figure S2A). Sequential salt extraction of chromatin-bound proteins further showed that Api5 binds more strongly to chromatin in malignant MCF10CA1a cells, eluting at higher salt concentrations (Figure 1F). Conversely, chromatin binding was weaker in the non-tumorigenic MCF10A and premalignant MCF10AT cells.

### Api5 deregulation causes genome instability

To investigate the phenotypic changes associated with Api5 depletion, we analysed nuclear volume and genomic content using 3D reconstruction and flow cytometry after DAPI staining, in MCF10CA1a cells with stable knockdown (KD) of Api5. Interestingly, cells with Api5 KD showed a significant reduction in nuclear volume and DNA content compared to empty vector (EV) cells (Figure 2A-C and G). In contrast, Api5 overexpressing (OE) cells exhibited increased nuclear volume and higher DNA content compared to EV cells (Figure 2D-F and H). Since nuclear volume change was detected in the cytochemistry analysis, changes should also be reflected in the flow cytometry analysis after G1 arrest with mimosine. Cell cycle arrest of MCF10CA1a EV and Api5 KD cells at the G1 phase using mimosine revealed an additional peak preceding the major G1 peak in Api5 KD cells, which was absent in EV cells. This additional G1 peak represents the cell population with low genomic DNA content (Figure S2B).

**Figure 2.**
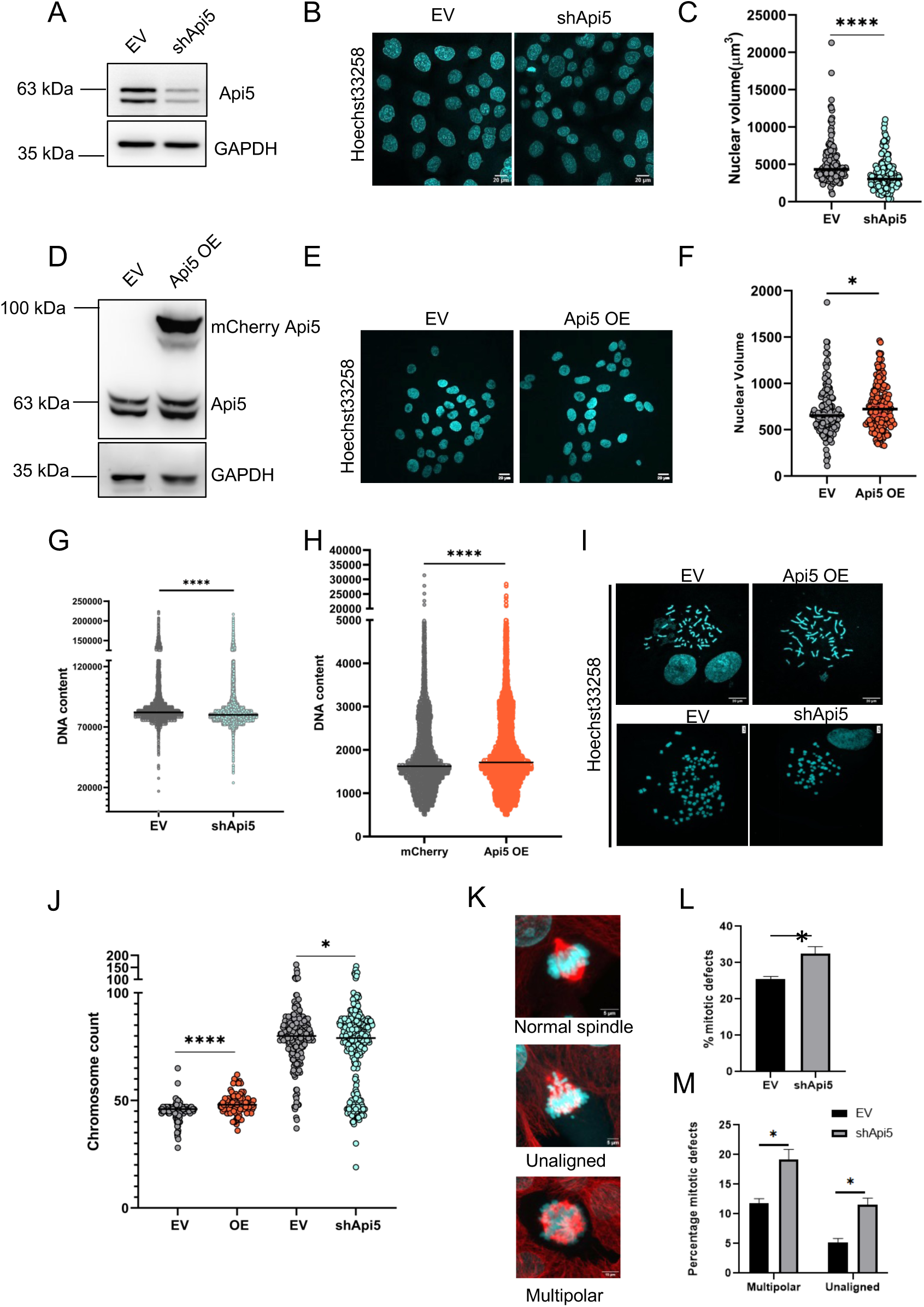
Api5 deregulation causes genomic instability. **A**. Western blotting was performed to confirm Api5 knockdown in MCF10CA1a cells. GAPDH was used as the loading control. **B**. Representative image showing reduction in nuclear size in MCF10CA1a shApi5 cells (scale bar 20μM). **C**. Nuclear volume analysis of MCF10CA1a EV and shApi5 cells using Imaris software (N = 3). ****P < 0.001. **D**. Api5 overexpression was confirmed by immunoblotting in stable cells generated using lentiviral-mediated overexpression of Api5 in MCF10A cells. GAPDH was used as the loading control. **E**. Representative image showing an increase in nuclear size in Api5 OE cells compared to EV cells (scale bar 20μM and 5μM). **F**. Nuclear volume analysis of MCF10A EV and Api5 OE cells using Imaris software (N = 3). ****P < 0.001. **G**. DNA content analysis was performed using flow cytometry in MCF10CA1a shApi5 cells after staining cells with DAPI; values represent median (N = 3). ****P < 0.001. **H**. DNA content analysis done by flow cytometry in MCF10A Api5 OE cells after staining cells with DAPI; values represent median (N = 3). ****P < 0.001. **I**. Chromosome spreads were performed in Api5 OE and Api5 KD cells (scale bar 20 μM). **J**. Chromosome counts presented as scatter plots, values represent the median (N = 3). Asterisks indicate statistically significant differences as determined by the Mann-Whitney test: *P < 0.05, ****P < 0.001. **K**. Representative images of mitotic defects analysis in the Api5 KD cells (scale bar 5μM). **L**. Total mitotic defects are quantified and represented as a bar graph (N = 4) *P < 0.05. **M**. Bar graph representing types of mitotic defects found upon Api5 KD in MCF10CA1a cells (N = 4) *P < 0.05.

To determine whether the observed changes in DNA content are associated with alterations in chromosome number, we performed metaphase chromosome spreads. Consistent with the DNA content measurements, cells with Api5 KD showed a significant reduction in chromosome number, whereas Api5 OE cells exhibited an increase in chromosome number compared with their respective controls (Figure 2I and J).

Since chromosomal gain or loss can arise from mitotic defects, we examined the mitotic defects in cells with altered Api5 expression. Api5 KD cells exhibited a significantly higher frequency of mitotic abnormalities, including multipolar spindle formation and chromosome misalignment, compared to EV controls (Figure 2K-M). Along with mitotic defects, MCF10CA1a cells with Api5 KD also exhibited a high incidence of micronuclei (Figure S2C and D), suggesting an association between Api5 KD and disrupted mitotic fidelity that may contribute to the observed changes in chromosome number.

To further test the clinical implications of Api5 in genomic instability, publicly available tumour datasets from The Cancer Genome Atlas (TCGA) were selected. Analysis of 2,858 breast invasive carcinoma samples revealed a positive correlation between Api5 mRNA expression and genomic instability, as assessed by fraction of genome altered (FGA) (Figure 3A) and tumour break load (TBL) (Figure S2E), supporting the association between Api5 levels and chromosomal alterations observed in cellular models. Apart from mRNA expression, putative copy-number alterations also positively correlated with genomic instability in breast invasive carcinoma datasets (Figure S2F).

**Figure 3.**
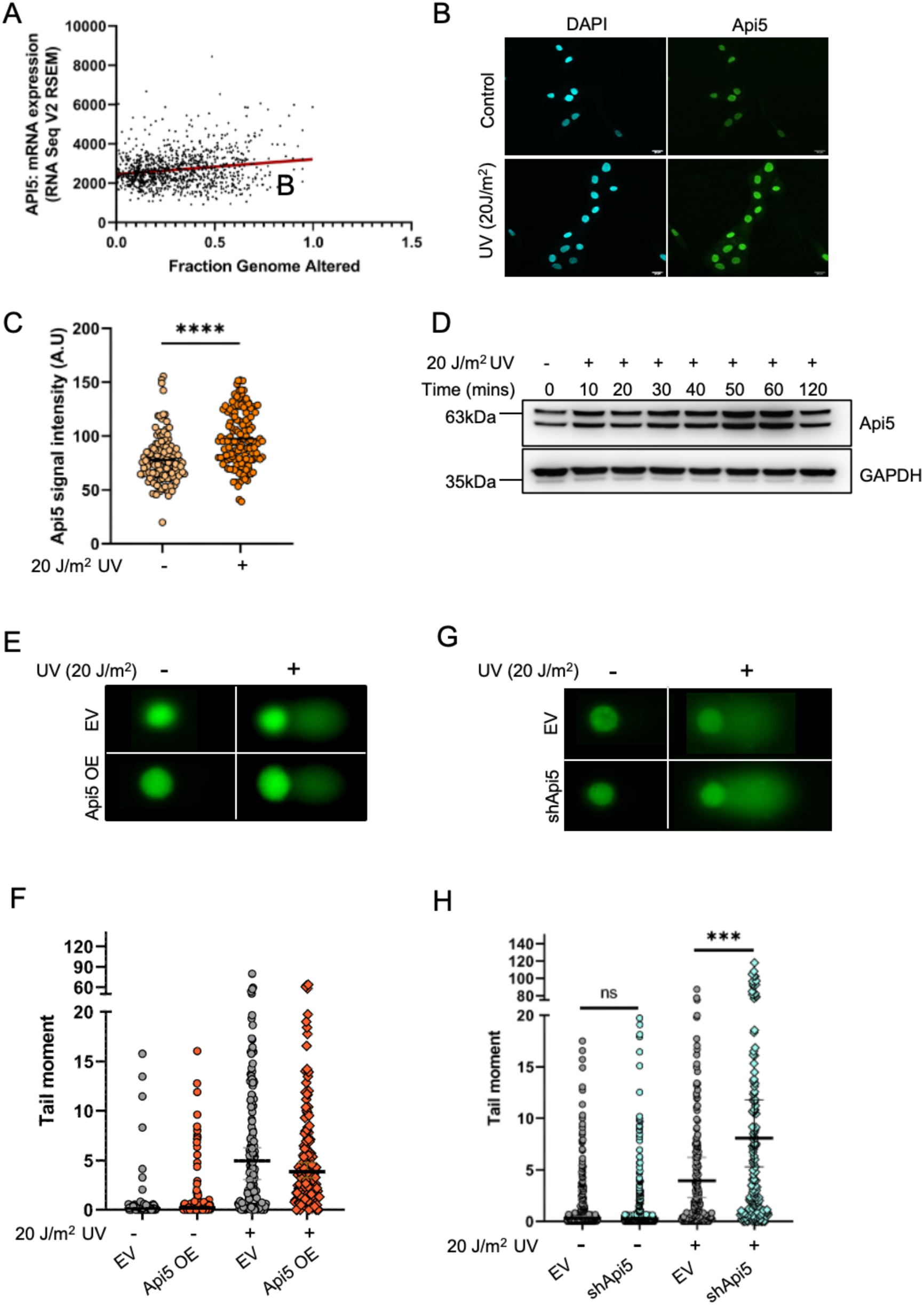
Api5 deregulation affects DNA damage. **A**. Scatter dot plot representing the breast invasive carcinoma datasets with fraction of genome altered on the x-axis and with mRNA expression levels quantified using RSEM on the y-axis. Regression line in black passing through the dots with ****P < 0.001 (n=1800). The results shown here are, in whole or in part, based on data generated by the TCGA Research Network. **B**. Immunocytochemistry representing the increase in Api5 levels following UV damage in MCF10CA1a cells. **C.** Quantification of signal intensity of Api5 from immunocytochemistry images after UV damage in MCF10CA1a (N=3, n≥800) ****P < 0.001. **D.** Western blot representing an increase in Api5 levels in lysates collected every 10 minutes after UV damage (N=3). **E**. Representative images of alkaline comet assay performed on MCF10A cells with Api5 OE and corresponding empty vector irradiated with 20 J/m^2^ UV dose. **F** Quantified values of comet tail moment in MCF10A EV and Api5 OE cells (N=3, n≥180) *P < 0.05. **G**. Representative Images of alkaline comet assay performed on MCF10CA1a cells with Api5 KD and corresponding empty vector irradiated with UV dose of 20 J/m^2^. **H**. Quantified values of comet tail moment in MCF10CA1a EV and shApi5 cells (N=3, n≥180) **P < 0.01.

### Api5 deregulation affects DNA damage sensitivity following UV exposure

Given the genomic instability phenotypes observed upon Api5 deregulation and previous reports linking Api5 to cisplatin resistance, we next examined the effect of UV-induced DNA damage on Api5 protein levels in cells. Immunoblot analysis revealed that Api5 levels progressively increased following 20 J/m^2^ UV damage, peaking at approximately 60 minutes post-treatment in MCF10CA1a (Figure 3B-D) and MCF10A cells (Figure S3A).

As Api5 levels increased after UV damage, we assessed the extent of DNA damage in Api5-depleted conditions using the comet assay. In the alkaline comet assay, which detects single-strand breaks (SSBs) and alkali-labile sites, Api5 KD cells exhibited significantly higher median comet tail moment values than EV cells, indicating increased DNA damage (Figure E and F). In contrast, Api5 OE cells showed a significant reduction in alkaline comet tail moment value relative to EV cells, suggesting decreased sensitivity to UV-induced DNA damage (Figure 3G and H).

In the neutral comet assay, which primarily detects double-strand breaks (DSBs), no significant differences in comet tail moment were observed between Api5 deregulated cells and their respective controls (Figure S3 B-E).

To further evaluate DNA damage, the expression of γH2AX, a marker for DNA double-strand breaks, was analysed. Following UV exposure, Api5 KD cells exhibited a significant increase in cells positive for γH2AX compared with EV controls (Figure 4A and B), whereas Api5 OE cells showed a marked reduction in γH2AX-positive cells relative to their corresponding EV controls (Figure 4C and D). Together, these results indicate that Api5 reduces the accumulation of UV-induced DNA damage and helps maintain genomic stability.

**Figure 4.**
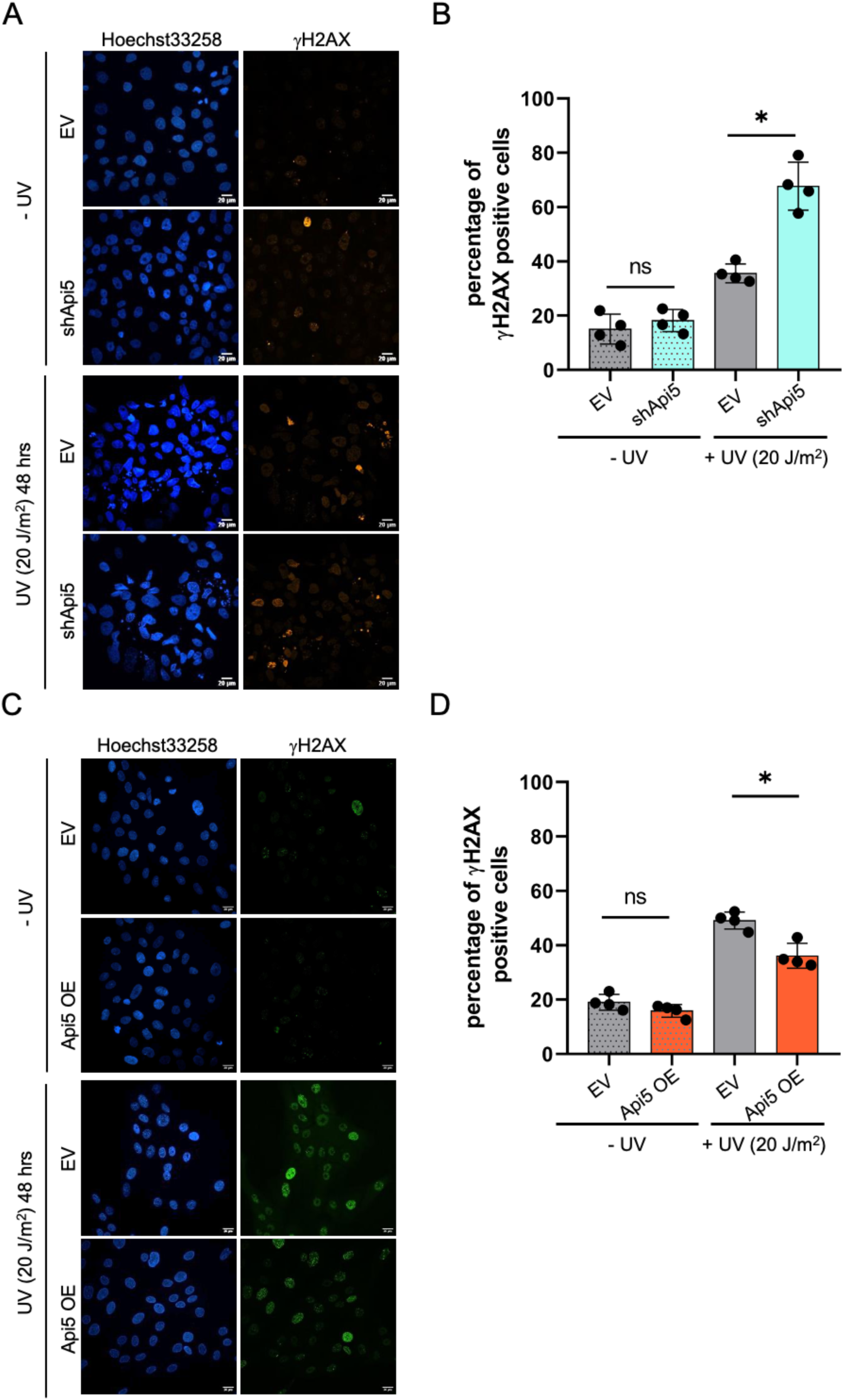
Api5 deregulation affects γH2AX accumulation after UV exposure. **A.** Representative image of γH2AX (orange) foci formation in MCF10CA1a EV and Api5 KD cells, 48 hours after exposure to 20 J/m^2^ UV and without UV exposure experimental control. Blue represents Hoechst 33258-stained nucleus. Scale bar: 20 µm. **B**. Quantification of the γH2AX positive cells in MCF10CA1a EV and Api5 KD, 48 hours after 20 J/m^2^ UV damage and without UV damage, showing an increase in γH2AX positive cells in Api5 KD cells. (N=3, n≥50). **C**. Representative images showing γH2AX (green) foci in MCF10A EV and Api5 OE cells, with or without exposure to 20 J/m² UV for 48 hours. Nuclei are stained blue with Hoechst 33258 dye. Scale bar:20 µm. **D.** Quantification of the γH2AX-positive cells in untreated and 20 J/m² UV-treated for 48 hours in MCF10A EV and Api5 OE cells (N=3, n≥50).

### Api5 deregulation affects the DNA Damage Response and repair pathways

To elucidate the role of Api5 in DNA damage repair following UV irradiation, the levels of cyclobutane pyrimidine dimers (CPDs), a major UV-induced DNA lesion repaired through the nucleotide excision repair (NER) pathway, were examined. Immediately after 20 J/m^2^ of UV damage, CPD levels in Api5 KD cells were comparable to those observed in the MCF10CA1a EV controls, indicating that Api5 KD does not affect the initial formation of UV-induced DNA damage. However, after 1 hour of post-UV recovery, Api5 KD cells exhibited significantly higher CPD levels than EV cells, suggesting disruption of UV-induced lesion repair (Figure 5A and B).

**Figure 5.**
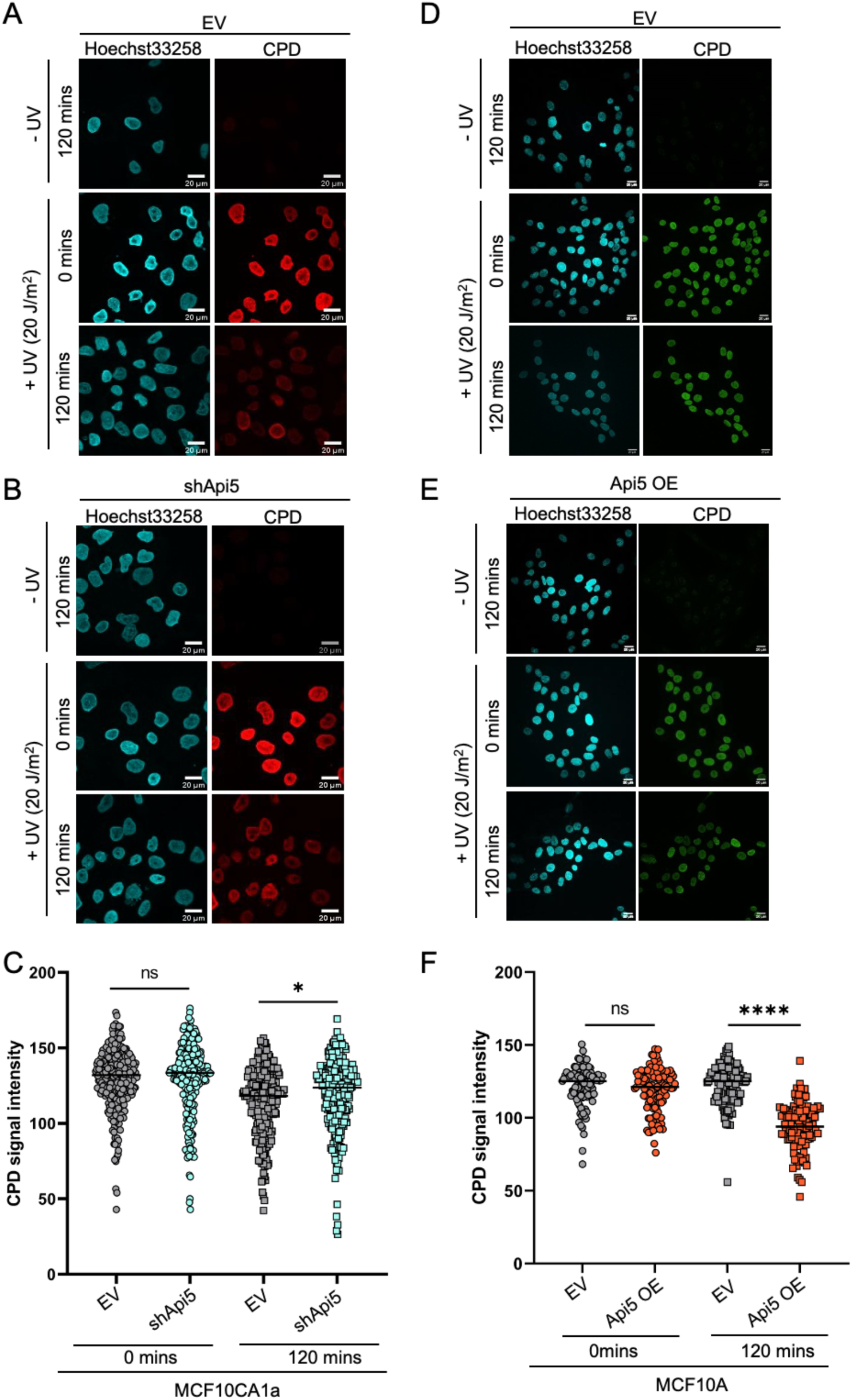
Api5 KD cells retain high levels of CPD after UV damage. **A**. and **B** CPD levels (red) were visualised in MCF10CA1a EV and Api5 KD cells that had been exposed to 20 J/m² UV damage for 0 and 120 mins. Nuclei were counter-stained with Hoechst 33258 (cyan). Scale bar: 20 µm. **C**. Quantification of CPD signal intensity from the EV and Api5 KD cells was shown as scatter plots with individual data points (N=3, n≥400). Mann–Whitney test, ****P < 0.001. **D** and **E**. CPD levels (green) in MCF10A EV and Api5 OE cells at 0 and 1 hour following 20 J/m² UV exposure. Hoechst 33258 was used to stain the nuclei (cyan). Scale bar: 20 µm. **E**. Quantification of CPD signal intensity from EV and Api5 OE cells shown as scatter plots with individual data points (N=3, n≥400). Mann-Whitney test, *P < 0.05.

To further corroborate this observation, similar experiments were performed in Api5 overexpressing MCF10A cells. In contrast to Api5 KD cells, Api5 OE cells displayed a significant reduction in CPD retention 1 hour after UV exposure compared with control cells, indicating improved repair of UV-induced DNA damage (Figure 5C and D). These findings suggest a potential role for Api5 in the nucleotide excision repair (NER) pathway.

### Api5 deregulation affects Chk1 phosphorylation

To determine whether the genomic instability phenotypes associated with Api5 deregulation arise from changes in the DNA damage response signalling pathway, the activation of the ATR signalling pathway was examined following UV-induced DNA damage in Api5-deregulated conditions. Cell lysates were collected 1 hour after UV irradiation, and the phosphorylation status of ATR pathway proteins was analysed. Api5 deregulation did not affect the phosphorylation of ATR at T1989, which is the phosphorylation indicative of ATR activation (Figure S4A-D). However, phosphorylation of Chk1 showed noticeable and consistent disruption. Api5 OE MCF10A cells showed increased Chk1 phosphorylation, whereas Api5 KD cells showed reduced Chk1 phosphorylation (Figure 6A-D).

**Figure 6.**
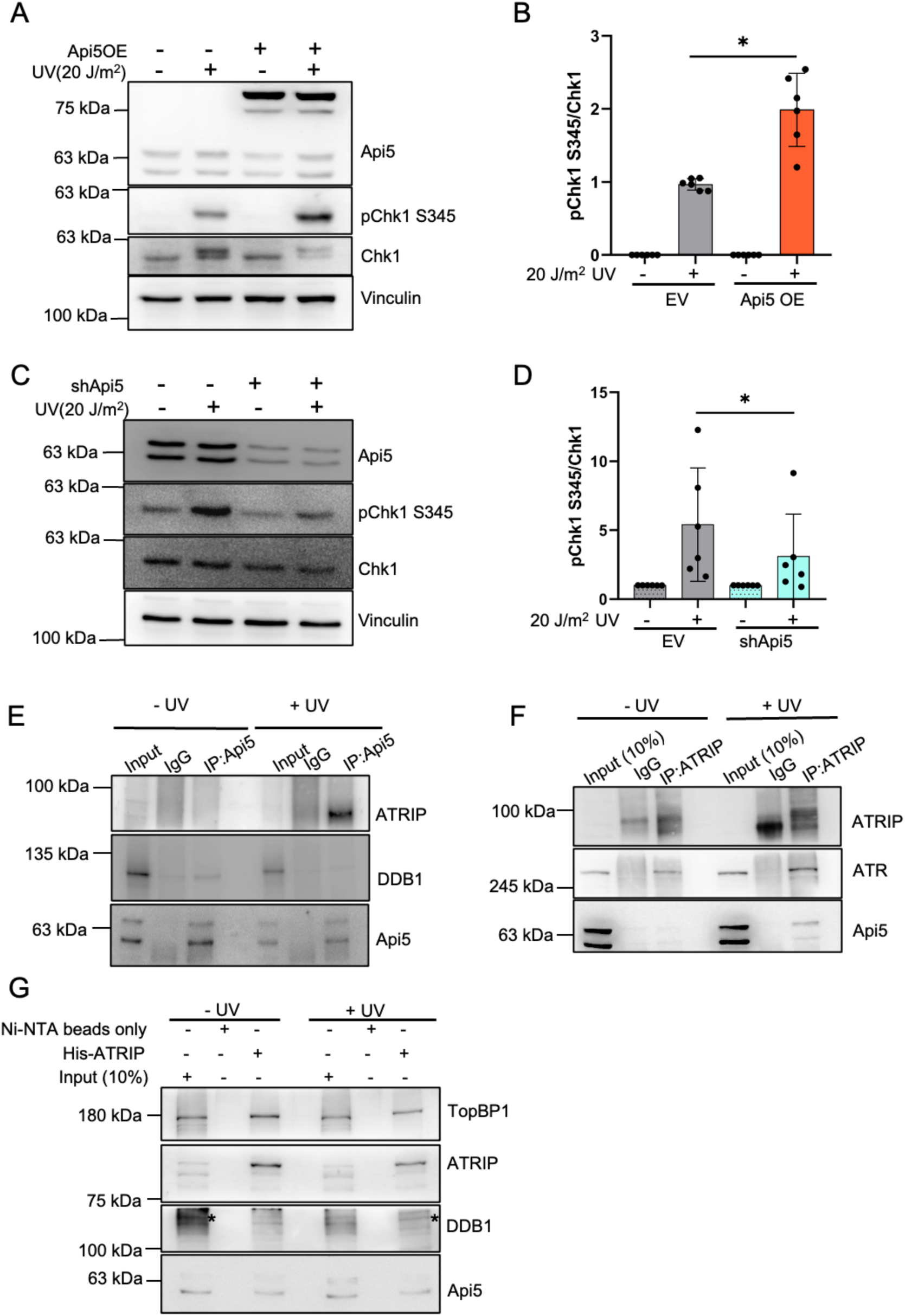
Phosphorylation of Chk1 at S345 is influenced by Api5 levels. **A**. Representative immunoblot of MCF10A EV and Api5 OE cells that had been exposed to 20 J/m^2^ UV-irradiation for 1 hour, showing increased Chk1 phosphorylation at S345 in UV-damaged Api5 OE cells (N=6), which was quantified using a Wilcoxon test (*P < 0.05) in panel **B**. **C**. Similar immunoblotting was performed in MCF10CA1a EV and shApi5 cells and probed for Chk1 phosphorylation at S345, in the absence and presence of 20J/m^2^ UV damage for 1 hour. **D**. Quantification of Chk1 phosphorylation in the MCF10CA1a EV and shApi5 cells showing a decrease in Chk1 activation in Api5 KD cells (N=6, Wilcoxon test *P < 0.05). **E**. Immunoprecipitation of Api5 was performed in MCF10A cells in the absence and presence of 20 J/m^2^ UV damage for 1 hr, and its interaction with DDB1 and ATRIP was studied (N=3). **F**. Immunoprecipitation of ATRIP from MCF10A cells in the absence and presence of 20 J/m^2^ UV damage for 1 hr, and its interaction with Api5 and ATR were evaluated (N=3). **G**. Ni-NTA bead-based pull-down of 6xHis ATRIP expressed in HEK293 cells shows the interaction of ATRIP with TopBP1, Api5, and DDB1 in the absence and presence of 20 J/m^2^UV damage for 1 hr.

To identify the interaction partners of Api5 that would explain the changes in Chk1 phosphorylation, co-immunoprecipitation (co-IP) was performed. The co-IP revealed that Api5 interacts with ATRIP following UV damage and with DDB1 in the absence of damage, suggesting an association with the nucleotide excision repair (NER) machinery and ATR pathway (Figure 6E). The Co-IP of ATRIP further confirmed its interaction with Api5 (Figure 6F). A pulldown assay using 6XHis-tagged ATRIP from cells transiently expressing ATRIP corroborated the interactions (Figure 6G). Taken together, these findings suggest that Api5 participates in the cellular response to UV-induced DNA damage by regulating checkpoint signalling and promoting DNA repair.

### Api5 deregulation affects cell survival after DNA damage and affects cisplatin resistance

As Api5 deregulation affects the DNA damage response pathway, it was essential to assess cell viability after genomic assault under Api5-deregulated conditions. The Api5 KD cells formed approximately 7% less colonies after UV exposure compared to EV (Figure 7A and B), and cisplatin treatment also showed similar results (Figure 7C and D). In the presence of Api5 overexpression, cells were more viable following UV treatment than in the EV condition, indicating resistance to DNA damage-induced cell death (Figure S5A and B). The Api5 OE cells exhibited 10% higher viability than the EV cells when treated with cisplatin (Figure S5C-E).

**Figure 7.**
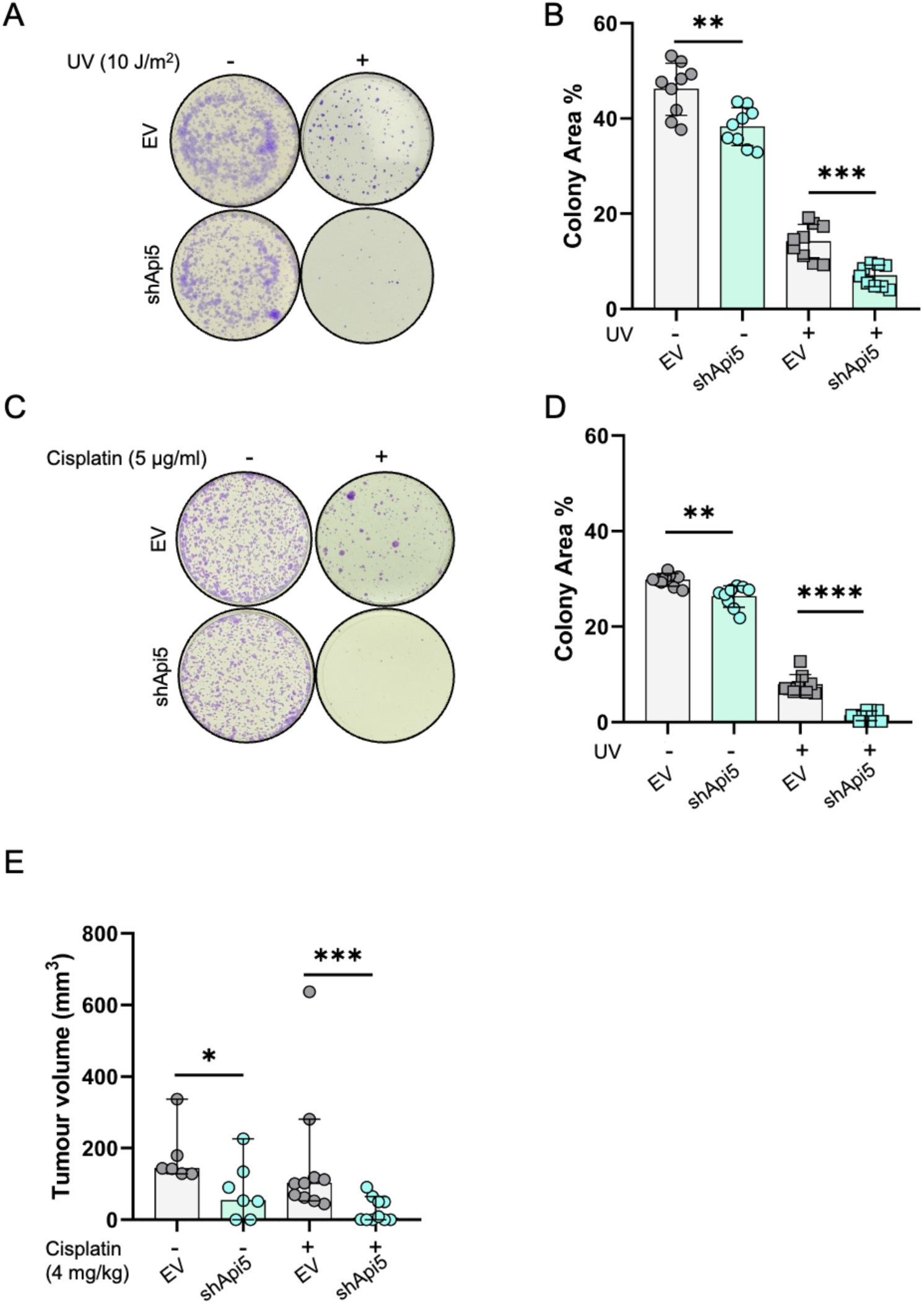
Api5 deregulation affects cell survival after DNA damage. **A**. Representative image from the colony formation assay of MCF10CA1a EV and shApi5 cells treated with UV light (10 J/m^2^). **B**. Quantification of the colony formation assay data. N=3, n=9, Mann Whitney test **P < 0.01, ****P < 0.001. **C**. Representative image from the colony formation assay done after cisplatin treatment in MCF10CA1a EV and shApi5 cells. **D**. Quantification of the colony area for cisplatin-treated and untreated cells. N=3, n=9, Mann-Whitney test **P < 0.01, ****P < 0.001. **E**. Quantification of the volumes of MCF10CA1a EV and shApi5 xenograft tumours induced by injection of MCF10CA1a shApi5 cells in the mammary fat pad, resected after cisplatin treatment. N≥6 for each group, Mann-Whitney test *P < 0.5, ***P < 0.001.

To evaluate whether these findings translate *in vivo*, xenograft studies were conducted in nude mice. Xenograft tumours were induced by injecting MCF10CA1a EV and shApi5 cells into the mammary fat pads of mice. Consistent with previous findings, Api5 knockdown significantly reduced the tumorigenic potential of the malignant MCF10CA1a cells. Furthermore, cisplatin (4 mg/kg) treatment to Api5 KD xenografts led to complete tumour regression in 40% of the mice, while the remaining 50% displayed a substantial reduction in tumour volume compared with the EV tumours treated with cisplatin (Figure 7E; Figure S6A-C). Taken together, these findings indicate that Api5 expression correlates with increased resistance to DNA damage and enhanced tumour cell survival.

## Discussion

Chemotherapy resistance associated with Api5-overexpressing tumours has been reported previously (Jang et al., 2017). However, studies have not investigated the role of the DNA damage response (DDR) pathways in Api5 deregulation. Many crucial proteins involved in DDR pathways interact with damaged DNA. The reports on chromatin association of Api5 (Garcia-Jove Navarro et al., 2013), its regulation of E2F1 target genes (Morris et al., 2006), and its interaction with the transcription factor ER-α (Basset et al., 2017) did not explain the relevance of Api5’s DNA binding, leaving this question unanswered. In this study, we identified a previously unrecognised DNA-binding property of Api5, along with differential association with chromatin during breast cancer progression, and its role in genomic instability. Our findings revealed that Api5 can directly bind to DNA via its C-terminal region, which harbours the DNA-binding domain. These observations suggest a potential role for Api5 in the chromatin-related processes.

Our findings identify Api5 as a regulator of genomic stability and a survival factor enabling cell viability under genotoxic stress. The reduction in nuclear size and DNA content under Api5-depleted conditions suggest chromosome loss rather than DNA condensation. This interpretation is further supported by the reduced DNA content and chromosome counts in Api5-depleted cells. Additionally, the increased prevalence of micronuclei in Api5-depleted conditions provide further evidence of Api5’s role in maintaining genomic integrity. The positive correlation of Api5 mRNA expression and the clinical parameters fraction of genome altered and tumour break load in the TCGA datasets highlights the clinical relevance of Api5 in genomic instability and cancer progression. Thus, the role of Api5 in genome maintenance is observable in the patient cancer datasets and cellular breast cancer progression models. The increase in the Api5 protein following UV damage indicates the functional relevance of maintaining Api5 levels for DNA damage repair. The increased sensitivity of Api5-depleted cells to DNA-damaging agents reinforces the relevance of Api5 in DNA repair and viability. The formation of CPD upon UV irradiation is an instantaneous photochemical process, and its complete removal requires at least 24 hours. Previous reports estimated peak CPD formation at 1-hour post-exposure. Here, it is interesting to note that malignant MCF10CA1a cells clear the CPD more efficiently than non-tumorigenic MCF10A cells, and Api5 OE certainly confers an advantage in repair, as MCF10CA1a is the cell line with the highest Api5 levels in the isogenic series. When the Api5 is knocked down in MCF10CA1a cells, CPD clearance is inefficient. As the NER pathway repairs the CPD, we can assume a role for Api5 in NER.

The retention of CPD in the Api5 KD cells and the significantly lower CPD levels in the Api5 OE cells indicate that Api5 is involved in DNA repair, not just viability. Smaller tumours and efficient regression after cisplatin treatment in Api5-depleted xenografts reproduced previous reports, albeit in the context of DNA repair and genomic stability. The increased viability of Api5 OE cells following DNA damage is a critical finding that could help explain reports of an association between high levels of Api5 and chemotherapy resistance. The use of multiple types of viability assessment under Api5-deregulated conditions strengthens the hypothesis with evidence.

The interaction of Api5 with DDB1 and ATRIP further demonstrates that Api5 is involved in the NER and DDR pathways, likely as a coordinator, as the complex forms even in the absence of DNA damage. However, we did not detect any interaction between Api5 and ATR or TopBP1 upon Api5 pull-down. Further analysis of the ATR pathway shows that ATR phosphorylation at T1989 does not change significantly, indicating that Api5’s role occurs after ATR activation. Even with no change in ATR activation in Api5-deregulated cells after UV exposure, Chk1 activation remains affected. It is important to note that Chk1 phosphorylation is considered a poor prognostic marker and is targeted in various cancers (Al-Kaabi et al., 2015; Alsubhi et al., 2016; Hwang et al., 2018; Zhang and Hunter, 2014). It is known that depletion of Chk1 leads to extensive DNA damage, as evidenced by increased γH2AX and RPA2 phosphorylation, which cause S-phase arrest, replication fork collapse, and failure to enter mitosis (Li et al., 2026; Michelena et al., 2019). The reduction in Chk1 phosphorylation at S345 observed in Api5 KD cells could explain genomic instability, including elevated γH2AX levels, prolonged comet tails, decreased viability, mitotic abnormalities, and heightened sensitivity to genotoxic stress. Using the MCF10 breast cancer progression model to assess DNA damage and repair, we have discovered a novel role for Api5 in genome stability. Moving forward, it will be interesting to investigate further how Api5 regulates Chk1 and its implications for its interactions with ATRIP and DDB1.

## Material and Methods

### Protein expression and purification

Api5 WT, Api5 Δ4 and Api5 Δ3 were earlier cloned into pGEX2TKcs using EcoRI (R3101S NEB) and NcoI (R3193S NEB) restriction enzymes and were used to transform BL21DE3 strains of *E. coli* using the heat shock method. The proteins were expressed by IPTG (4 mM) (67208 SRL) induction and incubation at 25°C for 12 hours with continuous agitation. The induced cells were collected by centrifugation at 4000 RCF for 15 minutes at 4 °C and lysed using a probe-based ultrasonicator (Sonics Vibra Cell^TM^) after re-suspending in extraction buffer (25 mM Tris-HCl (pH 8.0), 100 mM NaCl, 4% Triton X-100, and protease inhibitor cocktail (11836170001, Roche)). Lysate was clarified by centrifugation at 12000 RCF for 20 minutes at 4 °C. After clarification of the lysate, the protein was isolated by glutathione affinity purification beads (16101 Pierce™). Elution in 1xPBS was performed by bead thrombin digestion (1.12374, Sigma) to cleave off the glutathione S-transferase and release the Api5. Purified protein concentration was determined by the BCA assay (23225 Pierce™) and stored at -80 °C until use.

### Electrophoretic mobility shift assay (EMSA)

The purified protein and DNA (poly dA:dT 100-mer) were mixed with 10X binding buffer (100 mM Tris (pH 7.5 at 20 °C), 10 mM EDTA, 1 M KCl, 1 mM DTT, 50% vol/vol glycerol, 0.10 mg/mL BSA) and made up the volume to get 1X EMSA using dH_2_O. Api5 WT and Api5 Δ3 were mixed with varying concentrations of 0, 0.4, 0.8 and 1 μM with poly dA:dT 100-mer DNA in 0.3 μM in 1X binding buffer. The samples were run on a 1% agarose gel at 4°C in 0.25X TBE buffer at a constant voltage of 80 V. The gels were stained with SYBR™ Green (S9430, Sigma) and imaged with an Amersham™ Typhoon™ Laser Scanner.

### Cell lines and culture conditions

The MCF10A cell line was generously provided by Prof. Raymond C. Stevens (The Scripps Research Institute, USA), while the MCF10AT and MCF10CA1a cell lines were purchased from the Karmanos Cancer Institute, USA. Cells were regularly checked for mycoplasma contamination using Hoechst 33342 and the MAD™ Mycoplasma Detection Kit (OS-199305 S). MCF10A Api5 OE and MCF10CA1a shApi5 cells, along with their respective EVs, were obtained from a previously published study (Kuttanamkuzhi et al., 2023). The cells were cultured in High Glucose DMEM without sodium pyruvate (Gibco 11995081) supplemented with 5% horse serum (Gibco 16050122), 20 ng/mL EGF (Sigma-Aldrich E9644), 0.5 µg/mL hydrocortisone (Sigma-Aldrich H0888), 100 ng/mL cholera toxin (Sigma-Aldrich C8052), 10 µg/mL insulin (Sigma-Aldrich I1882-100MG), and 100 units/mL penicillin-streptomycin (Lonza 17-602E). During sub-culturing, cells were re-suspended in High Glucose DMEM without sodium pyruvate, containing 20% horse serum and 100 units/ml penicillin-streptomycin. HEK 293T cell line was a generous gift from Dr Jomon Joseph (National Centre for Cell Science, Pune, India). The MCF7 cell line was purchased from the European Collection of Authenticated Cell Cultures (ECACC). MCF7 cells were grown in High Glucose DMEM containing 10% foetal bovine serum (Invitrogen 10270106), 2 mM L-glutamine (Invitrogen 25030081) and 100 units/mL penicillin-streptomycin. HeLa cells were kindly provided by Dr Sorab Dalal (ACTREC, Navi Mumbai, India). HeLa cells were cultured in High Glucose DMEM (Invitrogen). This medium was supplemented with 10 % FBS, 100 units/mL penicillin, and 100 mg/mL streptomycin. All cells were maintained in 100 mm tissue culture-treated dishes (Invitrogen 150466, Thermo 150466) at 37°C in a humidified 5% CO_2_ incubator (New Brunswick Galaxy 170 R).

### Sequential salt extraction (SSE) of chromatin-bound protein

For sequential salt extraction of chromatin-bound proteins, the protocol standardised by (Porter et al., 2017) was followed, Briefly, 5 x 10^6^ cells were collected after trypsinisation and nuclei were isolated by mixing the cells with hypotonic solution (25 mM HEPES pH 7.6, 25 mM KCl, 5 mM MgCl_2_, 0.05 mM EDTA, 0.1% NP-40, and 10% glycerol and protein inhibitor cocktail) for 10 minutes at 4 °C on a rotor. The tubes were centrifuged at 6000 RCF for 5 minutes at 4 °C to separate the cytoplasmic and nuclear fractions. The nuclear pellet was re-suspended in 1X modified RIPA (mRIPA) (50 mM Tris pH 8.0, 1% NP-40 and PIC) and the proteins were eluted from the chromatin by pipetting exactly 15 times and incubating on ice for 3 minutes. After incubation, the tubes were centrifuged for 3 min at 6500 RCF at 4 °C. The supernatant was collected as the eluted protein fraction at 0 mM NaCl. The pellet was re-suspended in 100mM NaCl 1X mRIPA. The same procedure was followed for the sequential elution of chromatin-bound proteins from 100 mM to 500 mM NaCl in 1X mRIPA. The eluted protein fractions were analysed using immunoblotting.

### in silico analysis

The online DNA/RNA binding prediction bioinformatic tools DRNApred and DPbind were used to predict the DNA-binding region of Api5 (Hwang et al., 2007; Yan and Kurgan, 2017). AlphaFold structure prediction was used to model the folding of the Api5 deletion construct (Jumper et al., 2021). TCGA datasets with mRNA expression parameters from invasive breast carcinoma were downloaded. Both genomic instability indicators, i.e., the fraction of the genome altered and tumour break load, were analysed in relation to mRNA expression. Pearson correlation was used for statistical relevance.

### Plasmids and Cloning

AAC11 pGEX 4T11 and ATRIP pBSg constructs were a kind gift from Jean-Luc Poyet, INSERM, Paris, France and Lee Zou, Duke University, USA respectively. Full-length Api5 (1575bp) and its deletion constructs were cloned into pGEX 2TKcs for recombinant protein expression in *E. coli*. The API5 ORF was amplified using Phusion High-Fidelity DNA Polymerase using Api5 WT forward primer 5’-CATGATCCATGGATGCCGACAGTAGAGGAGCT-3’ and reverse primer 5’-CCAGAATTCTCAGTAGAGTCTTCCCCGAC-3’ containing EcoRI and NcoI restriction sites. For Api5 Δ3 cloning, a reverse primer with sequence 5’-ATCGGAATTCAAGTTGTCTGATATAAACTTGC-3’ was used along with Api5 WT forward primer. For Api5 Δ4 cloning, Api5 WT forward primer and reverse primer with sequence 5’-TAGCAGAATTCTCAGGGAGGGTTATAAAT-3’ were used.

### G1 arrest and Flow cytometry analysis

Cells that were 60% confluent were treated with 0.2 mM mimosine for 24 hours. The arrested cells were trypsinised and fixed with 70% ethanol. After fixation, the cells were stained with 50 µg/mL propidium iodide (PI) and analysed using BD Accuri™ C6 flow cytometer.

### Comet assay

The alkaline comet assay was performed using the previously reported protocol (Bodakuntla et al., 2014; Lu et al., 2017). The cells were treated with a UV-C dose of 10 J/m^2^ and incubated for 3 hours for the alkaline comet assay. After incubation, cells were harvested by trypsinisation, and 20000 cells suspended in PBS were mixed with 1% low-melting agarose and pipetted onto a glass slide coated with 1% low-melting agarose. The slides were then air-dried and immersed in alkaline lysis buffer (1.2 M NaCl, 100 mM EDTA, 10 mM Tris-HCl, pH >10, 0.1% sodium lauryl sarcosinate, 0.26 M NaOH) for lysis overnight at 4 ^°^C. The slides were electrophoresed after being rinsed in a pH >13 solution (0.03 M NaOH, 2 mM EDTA) for 25 minutes at 0.6 V/cm, with a distance between electrodes.

For the neutral comet assay for double-strand-break detection, neutral lysis solution (2% sarkosyl, 0.5 M Na_2_EDTA, 0.5 mg/ml proteinase K (pH 8.0)) was used to lyse the cells embedded in the low-melting agarose overnight at 37 ^°^C. The lysed slides were rinsed with neutral rinse and electrophoresis solution (90 mM Tris buffer, 90 mM boric acid, 2 mM Na_2_EDTA (pH 8.5)) and subjected to electrophoresis with similar conditions as those of the alkaline comet assay. The slides were washed by submerging in water for 5 minutes, followed by staining with 1X SYBR Green^TM^ in PBS for 20 minutes at room temperature in the dark. The excess stain was removed by rinsing in water, followed by imaging using a Leica DM6 epifluorescence microscope. The acquired images were analysed using Comet Score 2.0 software.

The comet tail moment is calculated using the following formula

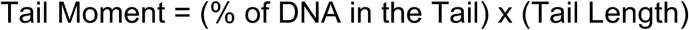

### Immunofluorescence and immunoblotting

The cells grown on glass coverslips were fixed with 4% PFA and permeabilised with 0.1% Triton X-100 in PBS, blocked with 5% FBS in PBS, then incubated overnight with the primary antibody diluted in blocking buffer, followed by a 1-hour incubation with the secondary antibodies also prepared in blocking buffer. The coverslips were mounted using Supernova mounting media and stored at 4^°^C until imaging. Imaging is performed using an Andor BC43 benchtop spinning-disk system for mitotic-defect slides and a Nikon AX confocal system for all other confocal images.

Api5 antibody (D-1) (sc-393341) was purchased from Santa Cruz Biotechnology. A 1:1000 dilution was used for western blotting, and a 1:50 dilution for immunofluorescence. Vinculin antibody (A2752) was purchased from ABclonal, and pATR T1989 (30632S) and ATR (13934S) were used at 1:10000 for western blotting at a dilution of 1:1000 and obtained from Cell Signalling Technology. Anti-H2AX phosphor Ser-139-FITC antibody (16-202A) was purchased from Millipore and was used for immunofluorescence at 1:100 dilution. Anti-cyclobutane pyrimidine dimer (CPD) monoclonal antibody (Clone TDM-2; CAC-NM-DND-001), used at a 1:1000 dilution for immunofluorescence, was purchased from Cosmo Bio USA. DDB1 polyclonal antibody (11380-1-AP) and Api5 polyclonal antibody (25689-1-AP) were obtained from Proteintech and used for western blotting at a 1:1000 dilution. GAPDH antibody (1:50000 dilution) (G9545-200UL) was purchased from Sigma. Goat anti-Rabbit IgG (H+L) Secondary Antibody, HRP (Thermo 31460) and anti-mouse HRP (Thermo 62-6520) were used for immunoblot at a 1:10000 dilution. Alexa fluor 488 goat anti-mouse (A11029 Thermo), Alexa fluor 488 goat anti-rabbit (A-11008 Invitrogen), Alexa fluor 568 goat anti-rabbit (Invitrogen A11011), and Alexa fluor 568 goat anti-mouse (Invitrogen A11004) were used as secondary antibodies for immunofluorescence at a 1:500 dilution.

### DNA content analysis

The DNA content analysis was performed by fixing cells at 70% confluence, trypsinising them, staining them with DAPI, and fixing them in ethanol prior to data acquisition using a BD FACSAria™. The acquired data are gated using early-passage MCF10A cells as a DNA-content baseline control, along with unstained controls for all sample groups.

### Co-immunoprecipitation

4 x 10^6^ cells were lysed using RIPA buffer (50 mM Tris, pH 7.4, 1% Triton X-100, and PIC), and protein quantification was performed using the BCA assay. 2 mg of bead-clarified lysates were incubated with 5 µg of antibody against the protein of interest and its respective isotype control overnight at 4°C on a rotor. After the incubation, the antibody was pulled down using 20 µl of protein A/G magnetic beads (Pierce^TM^ 88803) and washed with RIPA buffer (50 mM Tris, pH 7.4, 1% Triton X-100, and protein inhibitor cocktail). The washed beads were incubated in DTT-free Laemmli buffer and heated at 60 °C for 5 minutes, after which immunoblotting was performed.

### Pulldown assay

6xhis-tagged ATRIP was expressed in HEK293T cells by transient transfection using the pBSG01 plasmid. 48 hours after the transfection, the cells were treated with 20 J/m^2^ UV and lysed after 1 hour using RIPA buffer (50 mM Tris, pH 7.4, 1% Triton X-100, and protein inhibitor cocktail). The lysates were quantified, and 2 mg of lysate was mixed with 30 µl of Ni-NTA beads (Thermo R90115) for 2 hours at 4 °C on a rotor. The beads were washed 2 times with RIPA after incubation, and the bound proteins were mixed with Laemmli buffer and heated at 95 °C for 5 minutes, after which immunoblotting was performed.

### Cell viability assays

1.5 x 10^4^ cells were seeded in each well of a 96-well plate. Increasing concentrations of UV (10, 20, and 30 J/m^2^) and cisplatin (1-10 mg/ml) were tested, and samples were incubated for 24 hours. After the incubations, cell viability was assessed using a Calcein AM/PI-based cell viability assay kit (Olirum Scientific, OS-994-88) and BD FACSAria™ III Cell Sorter.

The Annexin-PI cell viability assay kit (Invitrogen™ V13242) was used to assess cell viability in MCF10A cells according to the manufacturer’s protocol, with a BD Accuri™ C6 flow cytometer.

For the PI exclusion assay, 1.5 x 10^4^ cells were seeded in each well in a 96-well plate. Various UV doses were tested on Api5 KD cells, and viability was assessed after trypsinisation and PI staining using a BD C6 flow cytometer.

### Colony formation assay

3 x 10^5^ cells were seeded in each well of 6-well plates and treated with UV-C (10 J/m^2^) and cisplatin (4 mg/ml). Immediately after UV exposure, the cells were trypsinised and 200 cells were seeded per well of a 6-well plate for a colony formation assay. Cisplatin treatment was for 24 hours prior to seeding for the colony formation assay. The cells were incubated until more than 50 cells per colony were present.

### Chromosome spreads

3 x 10^5^ cells were seeded in 6-well plates and enriched for the M-phase by thymidine-nocodazole treatment. The cells were treated with 2 mM thymidine for 20 hrs, followed by 10 hrs of nocodazole at 50 ng/ml. For chromosome preparation, cells were trypsinised, collected in 15 mL centrifuge tubes, and centrifuged at 122 x g for 5 min. The supernatant was discarded, leaving approximately 200 µL, and the pellet was gently resuspended. Cells were then treated with 5 mL of pre-warmed hypotonic solution (75 mM KCl) added slowly along the tube wall. Initially, 3 mL was added and mixed gently by inverting the tube, followed by the addition of 2 mL of hypotonic solution. Cells were incubated at 37 °C for 15 min. Subsequently, three drops of freshly prepared fixative solution (methanol: glacial acetic acid, 3:1) were added, gently mixed, and centrifuged at 122 x g for 5 min without brake. After discarding the supernatant, the cells were gently resuspended in 5 mL of fixative solution, which was added slowly along the tube wall while rotating the tube. Cells were stored overnight at 4 °C.

Glass slides were prepared by incubating them in 6 M HCl for at least 3 hrs at room temperature (RT), then washing under running tap water for 10 min, rinsing with distilled water, and air-drying. Slides were stored in 96% ethanol and dried with lint-free tissue immediately before use.

The following day, cells were centrifuged at 122 x g for 5 min without brake and washed three times with freshly prepared fixative solution (methanol: glacial acetic acid, 3:1). After the final wash, the pellet was resuspended in a sufficient volume of fixative to obtain an appropriate cell density. Approximately 20-30 µL of cell suspension was dropped onto clean, dry slides with the pipette positioned close to and parallel to the slide surface. Immediately after application, slides were exposed face-up to hot water steam (90°C) for 30 s to facilitate chromosome spreading. Prepared slides were examined under a phase-contrast microscope to assess cell density and distribution. The slides were then stained with DAPI, mounted with self-setting mounting medium (Olirum Scientific OS-062026S), and the number of chromosomes was counted using the ImageJ particle selection tool.

### *in-vivo* tumorigenicity assay

6 x 10^6^ cells were injected subcutaneously in the 4^th^ mammary fat pad of nude mice (B6.Cg-Foxn1nu /J, 6–8 weeks old) mixed with 1:1 diluted Matrigel^®^:PBS mixture. Tumour size was measured using a digital vernier calliper (Jiavarry). Cisplatin treatment (4 mg/kg) regime started when the tumours reached more than 5 mm in diameter. During cisplatin treatment, mice received 200 µl subcutaneous saline injections for hydration, Vitamin B (Procter & Gamble) and Vitamin E (Nukind Healthcare) supplements, and ThreptinTM biscuits (Raptakos, Brett & Co. Ltd) to prevent weight loss. The cisplatin treatment consisted of 5 doses administered every other day. Five days of rest were given before the mice were sacrificed, and the tumour was resected. The study is reported in accordance with the ARRIVE guidelines 2.0. The *in vivo* tumorigenicity studies were approved by the Institutional Animal Ethics Committee (IAEC) (IAEC/2025_01/03).

### Statistical analysis

All statistical analyses were performed using GraphPad Prism. Data are presented as mean ± standard error of the mean (SEM), unless otherwise specified. A P-value of < 0.05 was considered statistically significant.

For comparisons between two groups, the non-parametric Mann–Whitney U test was used. For experiments involving multiple conditions or independent samples, appropriate non-parametric tests were applied as indicated in the figure legends.

For western blot quantification, band intensities were analysed using the ImageJ densitometry tool, normalised to loading controls, and statistical significance was assessed using the Wilcoxon signed-rank test.

Correlation analysis between mRNA expression levels and the fraction of the genome altered (derived from The Cancer Genome Atlas invasive breast carcinoma datasets) was performed using the Pearson correlation coefficient (r) to assess the strength and direction of the association.

All experiments were performed with at least three independent biological replicates unless otherwise stated. Statistical details, including sample size (n) and exact p-values, are provided in the corresponding figure legends.

## Online supplemental material

Table S1 shows the DP-Bind results for Api5’s predicted DNA binding sequence. Figure S1 shows representative EMSA images using Api5 WT, Api5 Δ3, and Api5 Δ4 proteins. Figure S2 shows the increase in Api5 levels in the MCF10 series and genomic instability caused by Api5 deregulation in cellular and TCGA data. Figure S3 shows the increase in Api5 levels post-UV treatment in MCF10A cells and the neutral comet assay assessing double-strand breaks after UV exposure in Api5-deregulated cells. Figure S4 shows ATR phosphorylation under Api5-deregulated conditions, with and without UV exposure. Figure S5 shows the cell viability in Api5 OE cells treated with UV and cisplatin. Figure S6 shows the standardisation of cisplatin treatment regime, tumour volume and images of the resected tumours after treatment from xenograft studies.

## Acknowledgements

We are thankful to Prof. Manoj Mahimkar from the Advanced Centre for Treatment, Research and Education in Cancer (ACTREC), Mumbai, India, Dr Manas Kumar Santra from the National Centre for Cell Science (NCCS), Pune, India and Prof. Lee Zou, Duke Cancer Centre, Durham, North Carolina, USA, for their valuable suggestions. We also acknowledge the IISER Pune Microscopy Facility for access to equipment and infrastructure, the National Facility for Gene Function in Health and Disease (IISER, Pune, India) for access to the animal facility and support for experiments and the IISER Pune-BD FACS facility. We thank the members of the Lahiri laboratory for helpful discussions and comments.

## Author Contributions

BA: Conceptualisation, Investigation, Formal analysis, Methodology, Validation, Writing-original draft. ML: Conceptualisation, supervision, Funding Acquisition, Visualisation, Resources, Writing-review & editing. AU, KM, AM, DV and AD: Methodology, Validation.

## Conflict of Interest

The authors declare no competing or financial interests.

## Funding

This study is supported by a grant (BT/PR42717/BRB/10/1994/2021) from the Department of Biotechnology (DBT, Government of India) and, in part, by the Indian Institute of Science Education and Research, Pune, Core funding. BA was funded through the Council of Scientific and Industrial Research (CSIR)-JRF fellowship. Department of Science and Technology, the Indian Institute of Science Education and Research Pune, and the Department of Biotechnology.

## Notes

### Competing Interest Statement

The authors have declared no competing interest.

## References

Al-Kaabi, M., A. Alshareeda, D. Jerjees, A. Muftah, A. Green, N. Alsubhi, C. Nolan, S. Chan, E. Cornford, and S.J.B.j.o.c. Madhusudan. 2015. Checkpoint kinase1 (CHK1) is an important biomarker in breast cancer having a role in chemotherapy response. 112:901-911.

Alsubhi, N., F. Middleton, T.M. Abdel-Fatah, P. Stephens, R. Doherty, A. Arora, P.M. Moseley, S.Y. Chan, M.A. Aleskandarany, and A.R.J.M.O. Green. 2016. Chk1 phosphorylated at serine345 is a predictor of early local recurrence and radio-resistance in breast cancer. 10:213-223.

Basset, C., F. Bonnet-Magnaval, M.G.-J. Navarro, C. Touriol, M. Courtade, H. Prats, B. Garmy-Susini, and E.J.O. Lacazette. 2017. Api5 a new cofactor of estrogen receptor alpha involved in breast cancer outcome. 8:52511.

Basu, A., and S.J.J.o.n.a. Krishnamurthy. 2010. Cellular responses to Cisplatin-induced DNA damage. 2010:201367.

Bessette, D.C., E. Tilch, T. Seidens, M.C. Quinn, A.P. Wiegmans, W. Shi, S. Cocciardi, A. McCart-Reed, J.M. Saunus, and P.T.J.P.o. Simpson. 2015. Using the MCF10A/MCF10CA1a breast cancer progression cell line model to investigate the effect of active, mutant forms of EGFR in breast cancer development and treatment using gefitinib. 10:e0125232.

Bhatia, S., and M.J.J. Lahiri. 2026. Mammosphere Assay Reveals Api5-Induced Stemness in Non-Tumorigenic Breast Epithelial Cell Lines.e69999.

Bodakuntla, S., L.A. V, S. Sural, P. Trivedi, and M.J.B.c. Lahiri. 2014. N-nitroso-N-ethylurea activates DNA damage surveillance pathways and induces transformation in mammalian cells. 14:287.

Bousquet, G., J.-P. Feugeas, Y. Gu, C. Leboeuf, M. El Bouchtaoui, H. Lu, M. Espié, A. Janin, and M.J.O. Di Benedetto. 2019. High expression of apoptosis protein (Api-5) in chemoresistant triple-negative breast cancers: an innovative target. 10:6577.

Bouwman, P., and J.J.N.R.C. Jonkers. 2012. The effects of deregulated DNA damage signalling on cancer chemotherapy response and resistance. 12:587–598.

Cho, H., J.-Y. Chung, K.-H. Song, K.H. Noh, B.W. Kim, E.J. Chung, K. Ylaya, J.H. Kim, T.W. Kim, and S.M.J.B.c. Hewitt. 2014. Apoptosis inhibitor-5 overexpression is associated with tumor progression and poor prognosis in patients with cervical cancer. 14:545.

Egeblad, M., and Z.J.N.r.c. Werb. 2002. New functions for the matrix metalloproteinases in cancer progression. 2:161–174.

Garcia-Jove Navarro, M., C. Basset, T. Arcondéguy, C. Touriol, G. Perez, H. Prats, and E.J.P.o. Lacazette. 2013. Api5 contributes to E2F1 control of the G1/S cell cycle phase transition. 8:e71443.

Han, B.-G., K.H. Kim, S.J. Lee, K.-C. Jeong, J.-W. Cho, K.H. Noh, T.W. Kim, S.-J. Kim, H.-J. Yoon, and S.W.J.J.o.B.C. Suh. 2012. Helical repeat structure of apoptosis inhibitor 5 reveals protein-protein interaction modules. 287:10727-10737.

Hwang, B.-J., G. Adhikary, R.L. Eckert, and A.-L.J.O. Lu. 2018. Chk1 inhibition as a novel therapeutic strategy in melanoma. 9:30450.

Hwang, S., Z. Gou, and I.B.J.B. Kuznetsov. 2007. DP-Bind: a web server for sequence-based prediction of DNA-binding residues in DNA-binding proteins. 23:634–636.

Imre, G., J. Berthelet, J. Heering, S. Kehrloesser, I.M. Melzer, B.I. Lee, B. Thiede, V. Dötsch, and K.J.T.E.R. Rajalingam. 2017. Apoptosis inhibitor 5 is an endogenous inhibitor of caspase-2. 18:733-744.

Jang, H.S., S.R. Woo, K.-H. Song, H. Cho, D.B. Chay, S.-O. Hong, H.-J. Lee, S.J. Oh, J.-Y. Chung, J.-H.J.E. Kim, and M. Medicine. 2017. API5 induces cisplatin resistance through FGFR signaling in human cancer cells. 49:e374-e374.

Järvinen, A.K., R. Autio, S. Kilpinen, M. Saarela, I. Leivo, R. Grénman, A.A. Mäkitie, O.J.G. Monni, Chromosomes, and Cancer. 2008. High-resolution copy number and gene expression microarray analyses of head and neck squamous cell carcinoma cell lines of tongue and larynx. 47:500–509.

Jumper, J., R. Evans, A. Pritzel, T. Green, M. Figurnov, O. Ronneberger, K. Tunyasuvunakool, R. Bates, A. Žídek, and A.J.n. Potapenko. 2021. Highly accurate protein structure prediction with AlphaFold. 596:583–589.

Kim, Y.S., H.J. Park, J.H. Park, E.J. Hong, G.-Y. Jang, I.D. Jung, H.D. Han, S.-H. Lee, M.-C. Vo, and J.-J.J.O. Lee. 2018. A novel function of API5 (apoptosis inhibitor 5), TLR4-dependent activation of antigen presenting cells. 7:e1472187.

Koci, L., K. Chlebova, M. Hyzdalova, J. Hofmanova, M. Jira, P. Kysela, A. Kozubik, Z. Kala, and P.J.O.l. Krejci. 2012. Apoptosis inhibitor 5 (API-5; AAC-11; FIF) is upregulated in human carcinomas in vivo. 3:913-916.

Krejci, P., K. Pejchalova, B.E. Rosenbloom, F.P. Rosenfelt, E.L. Tran, H. Laurell, and W.R.J.J.o.L.B. Wilcox. 2007. The antiapoptotic protein Api5 and its partner, high molecular weight FGF2, are up-regulated in B cell chronic lymphoid leukemia. 82:1363-1364.

Kuttanamkuzhi, A., D. Panda, R. Malaviya, G. Gaidhani, and M.J.B.c. Lahiri. 2023. Altered expression of anti-apoptotic protein Api5 affects breast tumorigenesis. 23:374.

Li, S., D. Zhu, M. Tang, M. Huang, X. Feng, L. Nie, H. Zhang, L. Yin, S. Keast, C.J.C.D. Yang, and Disease. 2026. CHK1 is an integral regulator of DNA replication in human cells.

Lu, Y., Y. Liu, and C. Yang. 2017. Evaluating In Vitro DNA Damage Using Comet Assay. 1940-087X. e56450 pp.

Martin, L.P., T.C. Hamilton, and R.J.J.C.c.r. Schilder. 2008. Platinum resistance: the role of DNA repair pathways. 14:1291–1295.

Michelena, J., M. Gatti, F. Teloni, R. Imhof, and M.J.J.o.C.B. Altmeyer. 2019. Basal CHK1 activity safeguards its stability to maintain intrinsic S-phase checkpoint functions. 218:2865-2875.

Morris, E.J., W.A. Michaud, J.-Y. Ji, N.-S. Moon, J.W. Rocco, and N.J.J.P.g. Dyson. 2006. Functional identification of Api5 as a suppressor of E2F-dependent apoptosis in vivo. 2:e196.

Nygren, P.J.A.O. 2001. What is cancer chemotherapy? 40:166–174.

Ou, Y.-H., P.-H. Chung, T.-P. Sun, and S.-Y.J.M.b.o.t.c. Shieh. 2005. p53 C-terminal phosphorylation by CHK1 and CHK2 participates in the regulation of DNA-damage-induced C-terminal acetylation. 16:1684-1695.

Paulsen, R.D., and K.A.J.D.r. Cimprich. 2007. The ATR pathway: fine-tuning the fork. 6:953–966.

Pommier, Y., O. Sordet, S. Antony, R.L. Hayward, and K.W.J.O. Kohn. 2004. Apoptosis defects and chemotherapy resistance: molecular interaction maps and networks. 23:2934–2949.

Porter, E.G., K.E. Connelly, and E.C.J.J.o.v.e.J. Dykhuizen. 2017. Sequential salt extractions for the analysis of bulk chromatin binding properties of chromatin modifying complexes.55369.

Ray, A., K. Milum, A. Battu, G. Wani, and A.A.J.D.r. Wani. 2013. NER initiation factors, DDB2 and XPC, regulate UV radiation response by recruiting ATR and ATM kinases to DNA damage sites. 12:273-283.

Sasaki, H., S. Moriyama, H. Yukiue, Y. Kobayashi, Y. Nakashima, M. Kaji, I. Fukai, M. Kiriyama, Y. Yamakawa, and Y.J.L.C. Fujii. 2001. Expression of the antiapoptosis gene, AAC-11, as a prognosis marker in non-small cell lung cancer. 34:53-57.

Song, K.-H., S.-H. Kim, K.H. Noh, H.C. Bae, J.H. Kim, H.-J. Lee, J. Song, T.H. Kang, D.-W. Kim, and S.-J.J.B.r. Oh. 2015. Apoptosis inhibitor 5 increases metastasis via Erk-mediated MMP expression. 48:330.

Song, K., H. Cho, S. Kim, H. Lee, S. Oh, S. Woo, S. Hong, H. Jang, K.H. Noh, and C.J.O. Choi. 2017. API5 confers cancer stem cell-like properties through the FGF2-NANOG axis. 6:e285-e285.

Yan, J., and L.J.N.a.r. Kurgan. 2017. DRNApred, fast sequence-based method that accurately predicts and discriminates DNA-and RNA-binding residues. 45:e84-e84.

Yazinski, S.A., and L.J.A.r.o.g. Zou. 2016. Functions, regulation, and therapeutic implications of the ATR checkpoint pathway. 50:155–173.

Zhang, Y., and T.J.I.j.o.c. Hunter. 2014. Roles of Chk1 in cell biology and cancer therapy. 134:1013-1023.

