## Supplementary information for "Api5 Regulates Genomic Stability and Chemotherapy Resistance in Cancer"

**Table S1. DP-Bind results for Api5's predicted DNA binding sequence (showing only the positive sequences from the output for DNA binding).** Scores > 0.5: Generally, indicate a positive prediction (the higher the score, the higher the confidence that the residue physically interacts with that type of nucleic acid). Scores < 0.5: Generally, indicate a negative prediction (the residue is unlikely to bind).

| Position | Residue | Pred1 | Score1 | Pred2 | Score2 | Pred3 | Score3 | Pred4 | Consensus |
|---|---|---|---|---|---|---|---|---|---|
| 483 | P | 1 | 0.5 | 1 | 0.5305 | 0 | 0.6401 | 1 | NA |
| 484 | P | 1 | 0.7321 | 1 | 0.8591 | 1 | 0.5619 | 1 | 1 |
| 485 | S | 1 | 0.8386 | 1 | 0.7381 | 1 | 0.8982 | 1 | 1 |
| 486 | G | 1 | 0.9136 | 1 | 0.9215 | 1 | 0.8699 | 1 | 1 |
| 487 | K | 1 | 0.9584 | 1 | 0.9661 | 1 | 0.9705 | 1 | 1 |
| 488 | Y | 1 | 0.9114 | 1 | 0.935 | 1 | 0.9364 | 1 | 1 |
| 489 | S | 1 | 0.9498 | 1 | 0.8148 | 1 | 0.9039 | 1 | 1 |
| 490 | S | 1 | 0.8118 | 1 | 0.6912 | 1 | 0.7378 | 1 | 1 |
| 491 | N | 1 | 0.9051 | 1 | 0.9028 | 1 | 0.932 | 1 | 1 |
| 492 | L | 1 | 0.699 | 0 | 0.5177 | 1 | 0.563 | 1 | NA |
| 493 | G | 1 | 0.6547 | 1 | 0.6037 | 1 | 0.7267 | 1 | 1 |
| 494 | N | 1 | 0.8407 | 1 | 0.7053 | 1 | 0.9009 | 1 | 1 |
| 495 | F | 1 | 0.6535 | 1 | 0.5295 | 1 | 0.6598 | 1 | 1 |
| 496 | N | 0 | 0.5962 | 0 | 0.5641 | 1 | 0.5588 | 0 | NA |
| 497 | Y | 0 | 0.6469 | 0 | 0.5641 | 1 | 0.621 | 0 | NA |
| 498 | E | 0 | 0.6388 | 0 | 0.7745 | 0 | 0.7837 | 0 | 0 |
| 499 | Q | 1 | 0.5838 | 1 | 0.5927 | 1 | 0.6755 | 1 | 1 |
| 500 | R | 1 | 0.9131 | 1 | 0.8536 | 1 | 0.9219 | 1 | 1 |
| 501 | G | 1 | 0.7043 | 1 | 0.7233 | 1 | 0.8271 | 1 | 1 |
| 502 | A | 1 | 0.9194 | 1 | 0.8835 | 1 | 0.954 | 1 | 1 |
| 503 | F | 1 | 0.932 | 1 | 0.9101 | 1 | 0.9506 | 1 | 1 |
| 504 | R | 1 | 0.9553 | 1 | 0.9332 | 1 | 0.9354 | 1 | 1 |
| 505 | G | 1 | 0.8923 | 1 | 0.8091 | 1 | 0.8277 | 1 | 1 |
| 506 | S | 1 | 0.9244 | 1 | 0.8477 | 1 | 0.919 | 1 | 1 |
| 507 | R | 1 | 0.9813 | 1 | 0.9812 | 1 | 0.9258 | 1 | 1 |
| 508 | G | 1 | 0.9499 | 1 | 0.9102 | 1 | 0.8911 | 1 | 1 |
| 509 | G | 1 | 0.9589 | 1 | 0.9579 | 1 | 0.968 | 1 | 1 |
| 510 | R | 1 | 0.9892 | 1 | 0.9865 | 1 | 0.9818 | 1 | 1 |
| 511 | G | 1 | 0.9794 | 1 | 0.9683 | 1 | 0.9141 | 1 | 1 |
| 512 | W | 1 | 0.9814 | 1 | 0.9831 | 1 | 0.9799 | 1 | 1 |
| 513 | G | 1 | 0.9661 | 1 | 0.9588 | 1 | 0.8332 | 1 | 1 |
| 514 | T | 1 | 0.9549 | 1 | 0.9666 | 1 | 0.965 | 1 | 1 |
| 515 | R | 1 | 0.9788 | 1 | 0.9659 | 1 | 0.9393 | 1 | 1 |
| 516 | G | 1 | 0.7802 | 1 | 0.8324 | 1 | 0.806 | 1 | 1 |
| 517 | N | 1 | 0.883 | 1 | 0.9149 | 1 | 0.9249 | 1 | 1 |
| 518 | R | 1 | 0.8232 | 1 | 0.789 | 1 | 0.8542 | 1 | 1 |
| 519 | S | 1 | 0.8358 | 1 | 0.931 | 1 | 0.6976 | 1 | 1 |
| 520 | R | 1 | 0.9237 | 1 | 0.8479 | 1 | 0.8913 | 1 | 1 |
| 521 | G | 1 | 0.8314 | 1 | 0.8389 | 1 | 0.7314 | 1 | 1 |
| 522 | R | 1 | 0.7 | 1 | 0.785 | 1 | 0.6436 | 1 | 1 |
| 523 | L | 0 | 0.881 | 0 | 0.9227 | 0 | 0.7851 | 0 | 0 |
| 524 | Y | 1 | 0.6778 | 1 | 0.6205 | 0 | 0.5782 | 1 | NA |

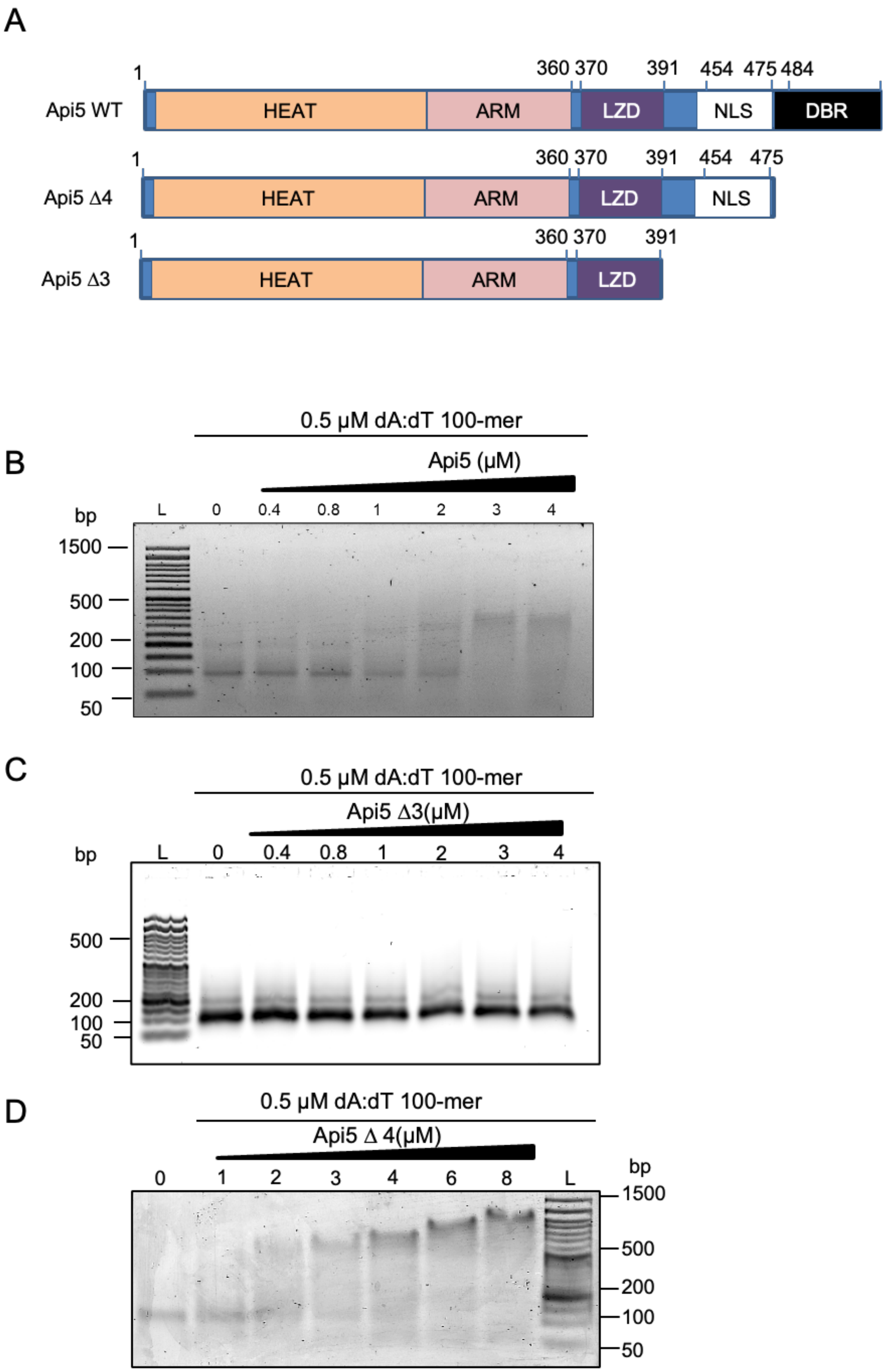


**Figure S1. DNA binding region of Api5 is at the C-terminal region. A**. EMSA of Api5 WT with poly dA:dT 100-mer was run on 1% agarose in 0.25X EMSA buffer. **B**.

EMSA of Api5 Δ3 with poly dA:dT 100-mer run on 1% agarose in 0.25X EMSA buffer. **C**. EMSA of Api5 Δ4 with poly dA:dT 100-mer run on 1% agarose in 0.25X EMSA buffer. N=3.

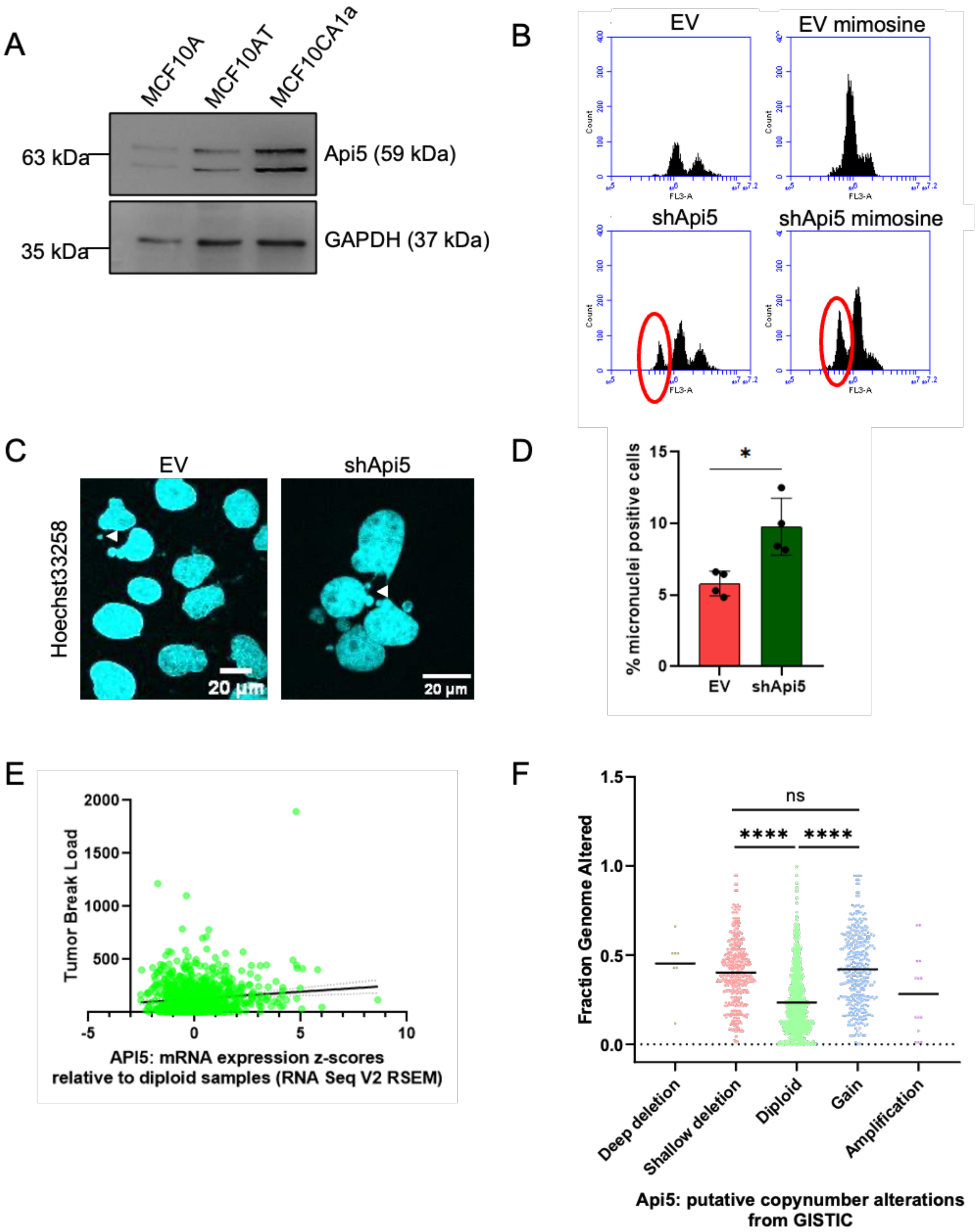


**Figure S2. Api5 deregulation causes genome instability**. **A.** Immunoblot showing Api5 levels in the MCF10 breast cancer progression series cells. **B**. Cytometry analysis of the MCF10CA1a EV and shApi5 cells treated with 0.2 mM mimosine for 24 hours and stained with DAPI, N=3. **C**. Representative image of micronuclei formation in MCF10CA1a EV and shApi5 cells stained with DAPI (cyan). **D**. Quantification of micronuclei in MCF10CA1a EV and shApi5 cells (N=3, n=150, Mann-Whitney; *P < 0.05). **E**. Scatter plot of tumour break load versus Api5 mRNA expression levels (RSEM) in breast invasive carcinoma samples (n = 1800). Each dot represents an individual sample, and the black line represents the linear regression fit. ****P < 0.001. Data were obtained, in whole or in part, from The Cancer Genome Atlas (TCGA) Research Network. **F**. Fraction of genome altered across different API5 copy number alteration groups obtained from GISTIC analysis, including deep deletion, shallow deletion, diploid, gain, and amplification. Samples with API5 gain exhibited a significantly higher fraction of the genome altered than diploid samples, whereas no significant difference was observed between gain and amplification groups. Statistical significance is indicated as **** (Mann-Whitney; P < 0.0001); ns, not significant.

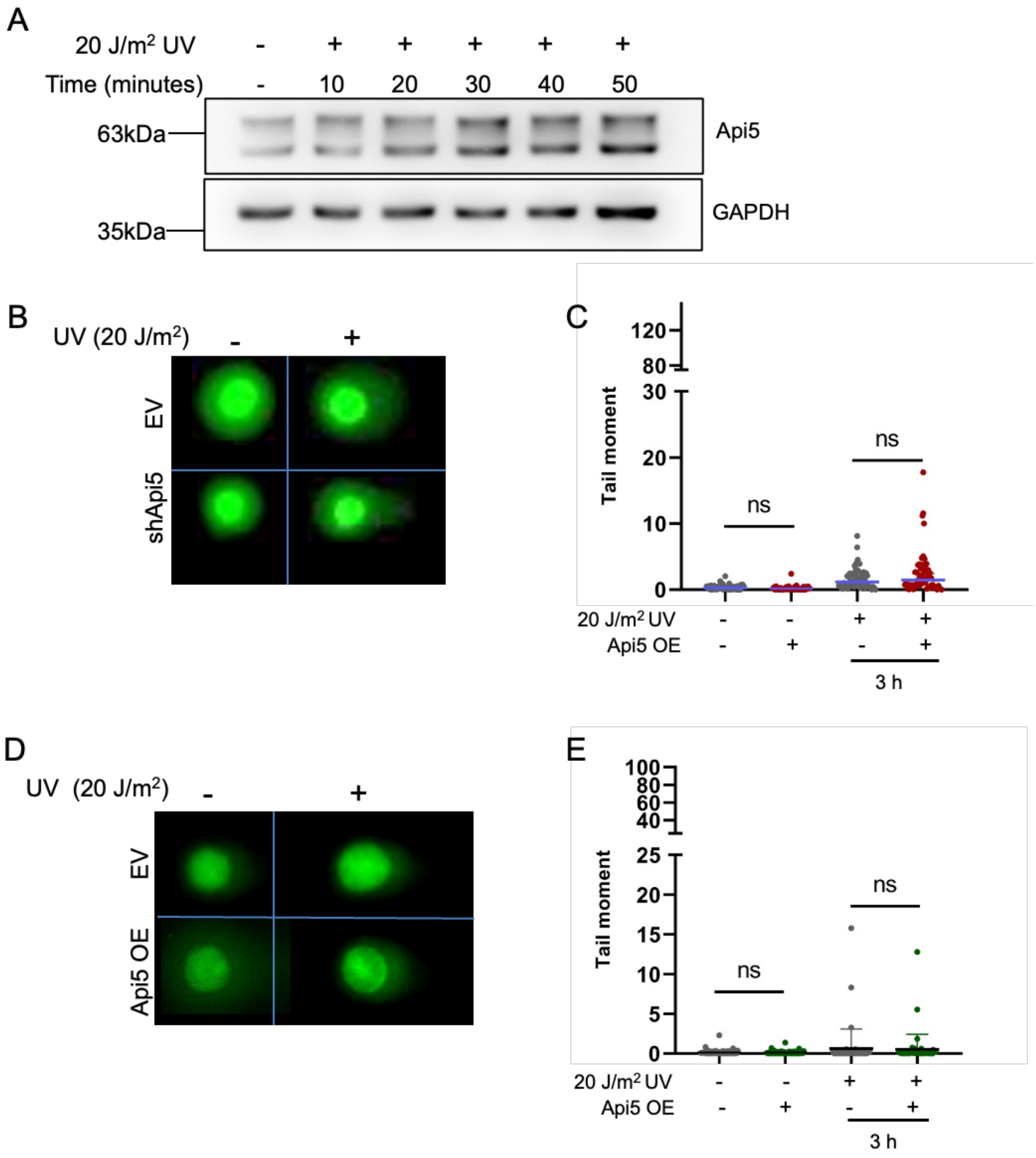


**Figure S3. Increase in Api5 levels after UV exposure and assessment of double-strand breaks in Api5 deregulated cells. A**. Representative western blot showing increased Api5 protein levels in MCF10A cells following UV-induced DNA damage. Cell lysates were collected at 10-minute intervals after UV exposure (N = 3). **B**. Representative images of the neutral comet assay performed on MCF10CA1a cells expressing either shApi5 or the corresponding empty vector (EV) following UV irradiation (20 J/m²). **C**. Quantification of comet tail moments in MCF10CA1a EV and shApi5 cells, presented as violin plots (N = 3, n ≤ 180 cells). *P < 0.05. **D**. Representative images of the neutral comet assay performed on MCF10A cells overexpressing Api5 (OE) or the corresponding empty vector (EV) following UV irradiation (20 J/m²). **E**. Quantification of comet tail moments in MCF10A EV and Api5 OE cells, presented as violin plots ($N = 3$, $n \leq 180$ cells). *$P < 0.05$.

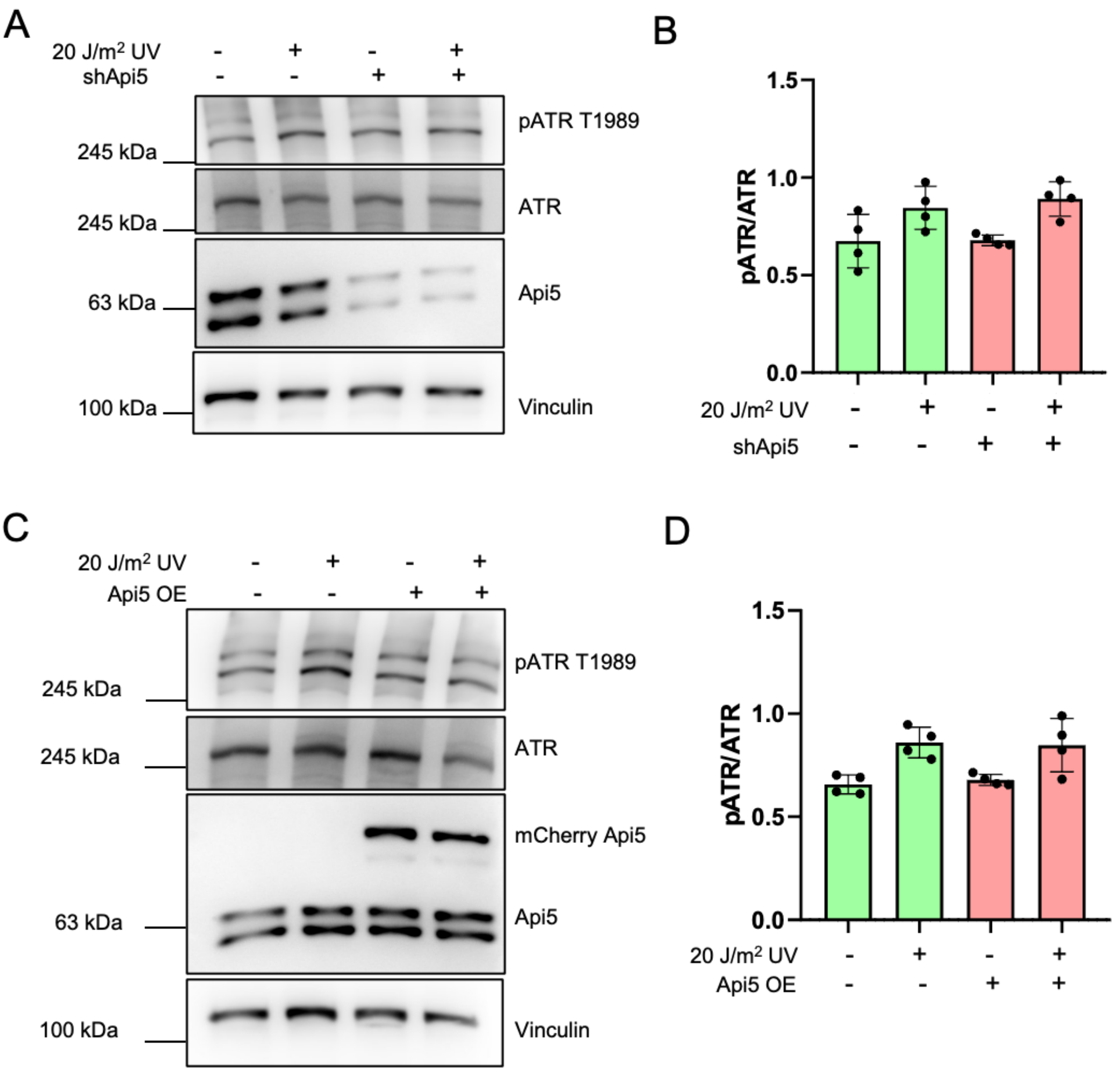


**Figure S4. ATR phosphorylation in Api5 deregulated cells following UV damage.** **A**. Representative Western blot showing ATR phosphorylation at Thr1989 in MCF10CA1a shApi5 cells, 1 hour after 20 J/m$^2$ UV irradiation (N = 4). **B**. Quantification of ATR phosphorylation at Thr1989 in MCF10CA1a shApi5 cells and MCF10A Api5-overexpressing (OE) cells. Data are presented from four independent experiments (N = 4). Statistical significance was determined using the Wilcoxon test. *P < 0.05. **C**. Representative Western blot showing Chk1 phosphorylation at Ser345 in MCF10A Api5-overexpressing (OE) cells 1 hour after UV irradiation (20 J/m²). **D**. Quantification of Chk1 phosphorylation at Ser345 in MCF10A Api5-overexpressing (OE) cells. Data are presented from six independent experiments (N = 6). Statistical significance was determined using the Wilcoxon test. *P < 0.05.

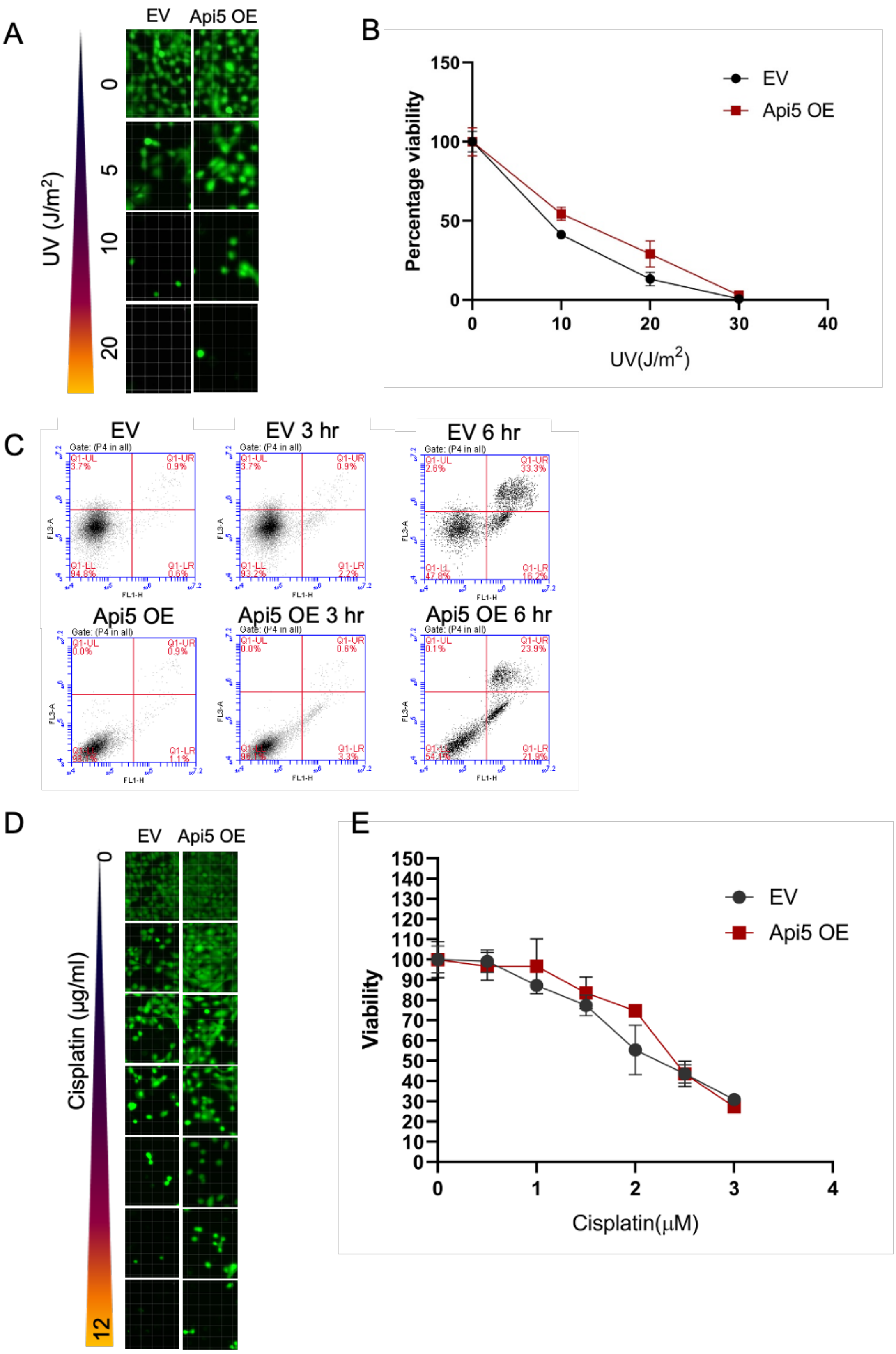


**Figure S5. Cell viability of Api5 OE cells following DNA damage. A.** Representative images of MCF10A Api5 OE cells and EV treated with increasing doses of UV after calcein AM-based cell viability assay. **B.** Fluorescence intensity measurement of calcein AM-based cell viability assay performed using MCF10A Api5 OE cells and EV treated with varying doses of UV. **C**. Annexin PI-based flow cytometry analysis of cell viability using cells treated with UV for 3 and 6 hours to identify early apoptotic cells. **D**. Representative images of MCF10A Api5 OE cells and EV treated with varying doses of cisplatin after calcein AM-based cell viability assay. **E**. Fluorescence intensity measurement of calcein AM-based cell viability assay performed using MCF10A Api5 OE cells and EV treated with increasing doses of cisplatin.

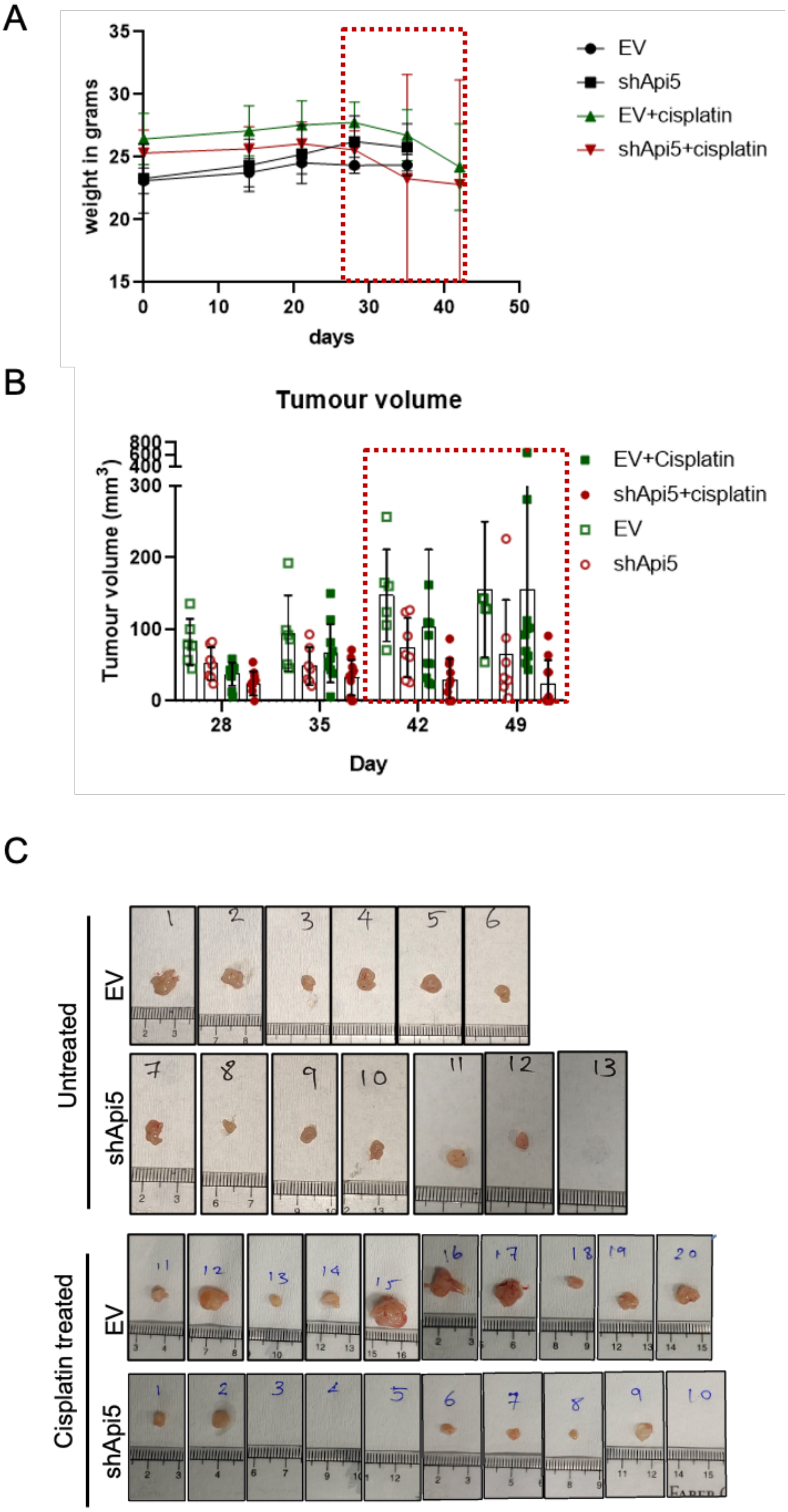


**Figure S6. Xenograft tumour volume and mice body weight changes during the treatment. A**. Body weight of mice from all experimental groups was recorded weekly throughout the study. The cisplatin treatment period is indicated by the red dashed rectangle. **B**. Tumour volume was measured weekly in all mice using a digital vernier calliper throughout the study. The cisplatin treatment period is indicated by the red dashed rectangle. **C**. Representative images of resected tumours from EV and shApi5 xenografts, grouped into untreated and cisplatin-treated cohorts. The animal identification number is shown above each tumour. **D**. Tumour volumes measured at the end of the experiment. Data were analysed using the Mann-Whitney test (N = 33). *P < 0.05, ****P < 0.001.
